# The carbapenem inoculum effect provides insight into the molecular mechanisms underlying carbapenem resistance in Enterobacterales

**DOI:** 10.1101/2023.05.23.541813

**Authors:** Alexis Jaramillo Cartagena, Kyra L. Taylor, Leslie C. Lopez, Jennifer Su, Joshua T. Smith, Abigail L. Manson, Jonathan D. Chen, Virginia M. Pierce, Ashlee M. Earl, Roby P. Bhattacharyya

## Abstract

Carbapenem-resistant Enterobacterales (CRE) are important pathogens that can develop resistance via multiple molecular mechanisms, including hydrolysis or reduced antibiotic influx. Identifying these mechanisms can improve pathogen surveillance, infection control, and patient care. We investigated, both phenomenologically and mechanistically, how resistance mechanisms influence the carbapenem inoculum effect (IE), a phenomenon where inoculum size affects antimicrobial susceptibility testing (AST). We demonstrated that any of seven different carbapenemases were sufficient to impart a meropenem IE when transformed into a laboratory strain of *Escherichia coli*. Across 106 clinical CRE isolates spanning 6 genera and 12 species, the carbapenem IE strictly depended on resistance mechanism: all 36 carbapenemase-producing CRE (CP-CRE) exhibited a clear IE, whereas 43 porin-deficient CRE displayed none. 27 isolates with both carbapenemase production and porin deficiency exhibited high-level resistance at all inocula, and still displayed an IE, albeit smaller in magnitude than CP-CRE with intact porins. Mechanistically, we found that clinical CP-CRE release carbapenemase activity into culture supernatant to protect other cells. Further, this released activity markedly increased upon exposure to lethal antibiotic doses, functionally consistent with altruism. Concerningly, 50% and 24% of CP-CRE isolates changed susceptibility classification to meropenem and ertapenem, respectively, across the allowable inoculum range in clinical testing guidelines. The meropenem IE, and the ratio of ertapenem to meropenem minimal inhibitory concentration (MIC) at standard inoculum, reliably identified CP-CRE. Understanding how resistance mechanisms affect AST could improve diagnosis and guide therapies for CRE infections.

**Importance:** Infections caused by carbapenem-resistant Enterobacterales (CRE) pose significant threats to patients and public health worldwide. Carbapenem resistance can occur through several molecular mechanisms, including enzymatic hydrolysis by carbapenemases and reduced influx via porin mutations. Identifying carbapenemase-producing isolates could enable tailored antibiotic selection to improve patient outcomes, and infection control measures to prevent further carbapenemase transmission. In a large collection of CRE isolates, we found that only carbapenemase-producing CRE exhibit an inoculum effect, in which their measured resistance varies markedly with cell density, which risks misdiagnosis. Further, this inoculum effect occurs under conditions where bacteria release carbapenemases to the community upon exposure to antibiotic that results in cell death, functionally consistent with altruism. Measuring this inoculum effect, or integrating other data from routine antimicrobial susceptibility testing, enhances carbapenem resistance detection, paving the way for more effective strategies to combat this growing public health crisis.

## Introduction

Carbapenems are a crucial class of β-lactam antibiotics with a broad spectrum of activity (1) due to their ability to withstand hydrolysis by many β-lactamases and their inhibition of cell wall synthesis across many bacterial species (2). The emergence of carbapenem-resistant Enterobacterales (CRE) poses a significant public health risk, with the US Centers for Disease Control and Prevention (CDC) designating them as an “urgent threat” due to their extensive drug resistance, high mortality rates, and treatment challenges (3, 4). The rapid global emergence of CRE has created an urgent clinical and epidemiological need to better understand, detect, and limit the spread of these pathogens (5).

CRE employ two major molecular mechanisms of carbapenem resistance: 1) expression of carbapenemases, which efficiently hydrolyze carbapenems, or 2) disruption of porins, which reduces influx of carbapenems into the periplasm where they act (6). This reduced periplasmic influx can potentiate β-lactamases with weak carbapenemase activity such as AmpC, CTX-M, SHV, TEM, or OXA-10, which together cause resistance (1, 7). Efflux pumps can also contribute to carbapenem resistance (8), though their role has been better characterized in non-Enterobacterales species like *Acinetobacter baumannii* and *Pseudomonas aeruginosa* (9, 10). Carbapenemase-producing CRE (CP-CRE) are of particular epidemiological concern due to their propensity to cause outbreaks (4, 11) and the risk of horizontal transfer of carbapenemase genes (12–14). Porin deficient CRE (PD-CRE) often exhibit impaired fitness (15) compared to CP-CRE because porins play important roles in nutrient import (16), osmotic homeostasis (15, 17), and membrane lipid asymmetry (18). Determining resistance mechanisms could lead to better infection control measures, such as patient isolation to prevent CP-CRE spread in hospitals (4, 13, 19). Many laboratories in high-resource settings now test for resistance mechanisms in CRE (20–22), but this requires additional time, equipment, expertise, and expenses. Capitalizing on existing antimicrobial susceptibility testing (AST) workflows to predict CRE resistance mechanisms and enhance efficiency of molecular testing deployment would represent a clinical advance.

Standardized AST methods and interpretive criteria are crucial to determine antimicrobial susceptibility of pathogens. Organizations like the Clinical and Laboratory Standards Institute (CLSI) classify pathogen susceptibility based on minimum inhibitory concentration (MIC) breakpoints (23–25) that are established and periodically revised (26) based on microbiological, pharmacokinetic-pharmacodynamic, and clinical data. While these breakpoints are intended to predict clinical response to therapy, not resistance mechanism, carbapenem MICs have been repurposed to prioritize isolates for molecular carbapenemase testing (25, 27). In the United States, only isolates that test phenotypically carbapenem resistant are typically tested for carbapenemases (24). Yet despite recently updated breakpoints (26), some carbapenemase producing isolates can still be misclassified as susceptible to carbapenems (28).

The inoculum effect (IE) is a long-recognized phenomenon whereby the measured MIC varies depending on the precise bacterial inoculum used in AST assays, which can lead to susceptibility misclassification and treatment failures (29–31). Measured MICs can vary markedly because of the IE, with high inocula withstanding antibiotic concentrations hundreds of fold higher than low inocula, consistent with the majority of resistance emerging from the community, rather than individual bacteria. Multiple factors are proposed to contribute to the IE, including metabolic state of the bacteria (32), growth productivity (33), antibiotic-target ratio (34), heteroresistance frequency (35), target proteolysis (36), and production of enzymes that inactivate antibiotics (37–41). The IE is frequently observed for some antibiotic classes including β-lactams (31, 42) but not commonly for others such as aminoglycosides. In *Staphylococcus aureus*, where the cefazolin IE has been thoroughly studied, β-lactamase carriage appears to be necessary but not sufficient to predict an IE (43–45). While the importance of the IE in clinical infections remains uncertain (46, 47), it has important implications for diagnostic accuracy. To minimize the impact of the IE, CLSI recommends a standard inoculum of 5 × 10^5^ colony forming units (CFU)/mL, with an acceptable range of 2 × 10^5^ to 8 × 10^5^ CFU/mL, for broth microdilution, the gold standard AST method (24). Even within this range, however, carbapenem susceptibility classification of Enterobacterales isolates may change due to the IE, underscoring the need for continued efforts to optimize AST methods (48). Thus, clarity on the mechanistic basis for the IE in the Enterobacterales has diagnostic and potentially therapeutic implications.

Here, we sought to systematically investigate how different genotypes conferring carbapenem resistance influence the IE and how understanding these relationships might be leveraged to improve clinical AST. We found that expression of distinct carbapenemases was sufficient to cause a marked meropenem IE in a laboratory strain of *Escherichia coli*. Further, in a collection of 106 clinical CRE isolates across 12 species of Enterobacterales (12, 14, 49), we observed that the carbapenem IE was fully predicted by resistance mechanism. Specifically, all CP-CRE exhibited a strong carbapenem IE, whereas all PD-CRE exhibited constant carbapenem MICs across a wide range of inocula. Isolates with both a carbapenemase and porin deficiency generally exhibited an IE and higher MICs than strains with either mechanism alone. Release of carbapenemase activity from clinical CP-CRE into the supernatant to confer transferrable protection to other cells was markedly enhanced upon exposure to lethal antibiotic doses, providing a plausible mechanistic basis for community protection, functionally consistent with altruism from antibiotic-mediated cell death. The carbapenem MICs of CP-CRE varied considerably, with a large proportion changing AST classification across the CLSI-acceptable inoculum range. Collectively, our results demonstrated that the IE distinguished CRE by resistance mechanism, better than either meropenem or ertapenem MIC alone. These findings suggested two phenotypic measures that predict carbapenem resistance mechanisms better than the MIC cutoffs currently used to prioritize molecular carbapenemase testing.

## Results

### Carbapenemases belonging to all four Ambler classes impart an inoculum effect

Carbapenemases are commonly divided into four classes based on protein sequence and enzymatic features (50, 51). To investigate if each carbapenemase class is sufficient to confer an IE, we systematically cloned seven different carbapenemase genes, each with its native promoter, into a low-copy pBAD33 plasmid (52): *bla*_KPC-3_ and *bla*_SME-2_ from Ambler class A; *bla*_IMP-4_, *bla*_NDM-1_, and *bla*_VIM-27_ from class B; *bla*_CMY-10_ (53) from class C; and *bla*_OXA-48_ from class D (Supplementary Table S1). We transformed each carbapenemase-encoding plasmid into *E. coli* K-12 MG1655, a carbapenem-susceptible laboratory strain, then conducted meropenem broth microdilution MIC assays across a broad range of inocula, comprising two-fold dilutions from 1.3 × 10^7^ to 1.6 × 10^3^ CFU/mL. As inoculum increased, all transformants showed a significant increase in MIC, while the MIC of the empty vector control did not (Fig. 1*A*). Most transformants had stable meropenem MICs from ∼10^3^ to ∼10^4^ CFU/mL, and all exhibited a strong linear increase in MIC across the ∼10^5^ to ∼10^7^ CFU/mL inoculum range, which includes the recommended inoculum range for clinical testing (Fig. 1*B*). In this strain background, NDM-1 (orange), SME-2 (pink), and IMP-4 (brown) imparted the highest levels of resistance, followed by VIM-27 (yellow). KPC-3 (teal) conferred high-level resistance at high inocula, but within the CLSI inoculum range (dashed lines in Fig. 1*B*) MICs remained within the intermediate classification (>1 µg/mL and <4 µg/mL). OXA-48 (violet) and CMY-10 (green) conferred the lowest meropenem MICs, which remained below 1 µg/mL until the highest inoculum tested and well within the susceptible classification in the CLSI range. The extended-spectrum beta-lactamase (ESBL) gene *bla*_CTX-M-15_, which often contributes to a CRE phenotype when paired with porin deficiency (54, 55), conferred only a minimal IE, similar to empty vector (Supplementary Fig. S1).

**Fig. 1.**
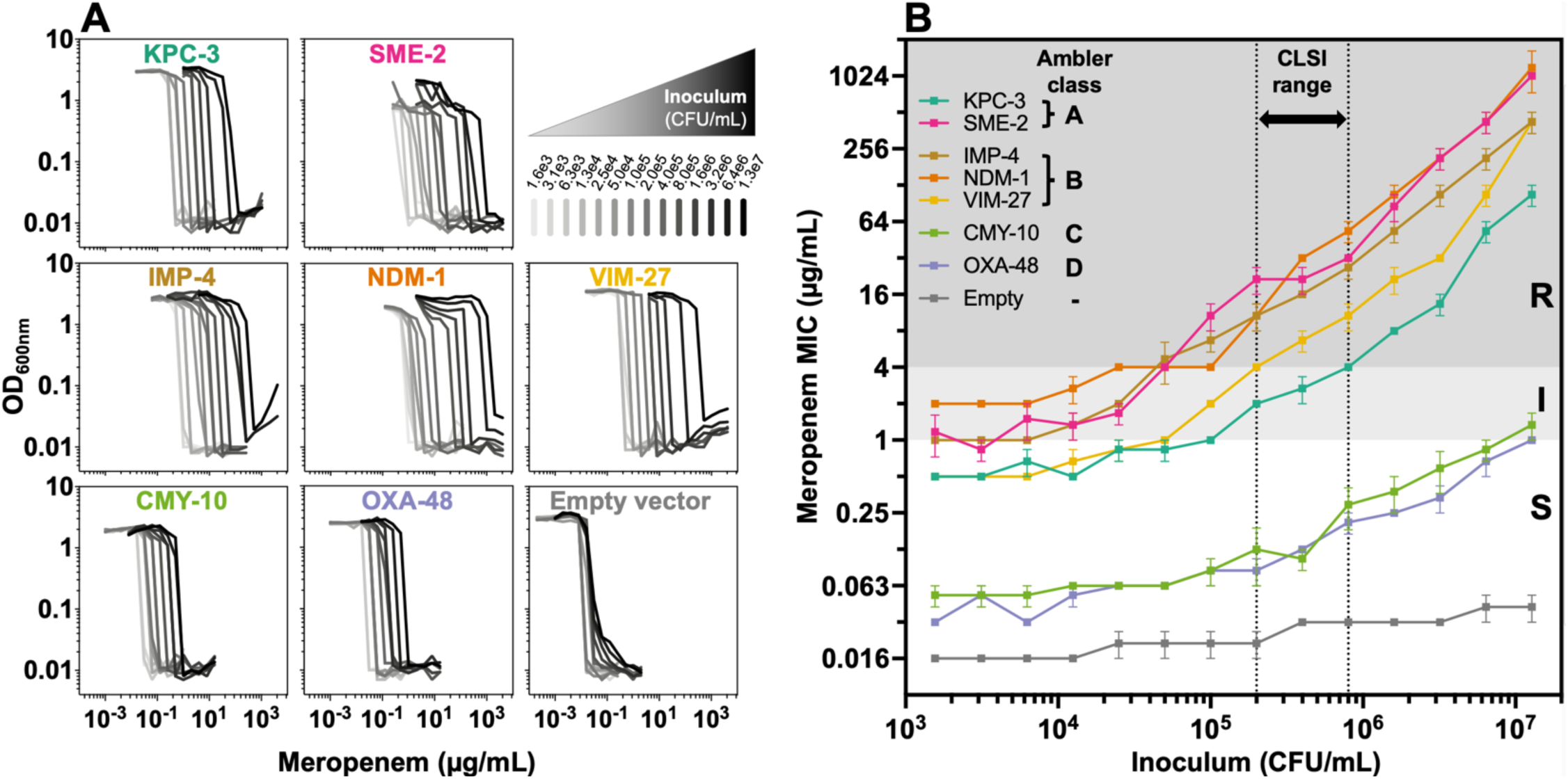
All carbapenemases impart a meropenem inoculum effect when expressed in *E. coli K12*. **(A)** Plots of optical density (OD) at 600 nm (y-axis) after overnight incubation with varying meropenem concentrations (x-axis) across a broad range of starting inocula (grayscale per top right panel) for *E. coli* K-12 transformed with each indicated carbapenemase. **(B)** Meropenem minimum inhibitory concentration (MIC) as a function of bacterial inoculum, colored by transformed carbapenemase. Each MIC point is the mean of three replicates (error bars = standard error of the mean). Vertical dotted lines represent the CLSI-acceptable inoculum range (2 to 8 x 10^5^ CFU/mL). The background is shaded by CLSI meropenem susceptibility breakpoints (S = susceptible, white; I = intermediate, light gray; R = resistant, gray).

### Inoculum effect of clinical CRE isolates depends on resistance mechanism

After establishing that all carbapenemases imparted a meropenem IE in *E. coli* K-12, we investigated the IE in clinical isolates harboring carbapenemases, porin disruptions, or both. We analyzed 106 clinical CRE isolates comprising 6 genera and 12 species (Supplementary Table S2), which we assigned to three groups based on their genotypes: CP-CRE, PD-CRE, and CP-CRE with porin deficiency (Fig. 2 and Supplementary Fig. S2, Table S2 and Table S3). We observed that all 36 CP-CRE harboring carbapenemases from one of six major families (KPC, NDM, IMP, VIM, OXA-48, and SME) displayed a meropenem IE similar to the *E. coli* K-12 carbapenemase transformants (Fig. 2*A* and Supplementary Fig. S2*A*). In contrast, 43 PD-CRE, many of which encoded ESBLs, exhibited a strikingly different pattern of resistance, with nearly constant MICs across the tested inoculum range (Fig. 2*B* and Supplementary Fig. S2*B*). The 27 isolates with both carbapenemase production and porin deficiencies, which we termed “hyper-CRE”, exhibited an IE and higher-level resistance across all tested inocula than isolates containing only one mechanism of resistance (Fig. 2*C* and Supplementary Fig. S2*C*). 5 ESBL-encoding meropenem-susceptible isolates exhibited no meropenem IE, nor did 5 other randomly selected meropenem-susceptible isolates (Supplementary Fig. S2*D*).

**Fig. 2.**
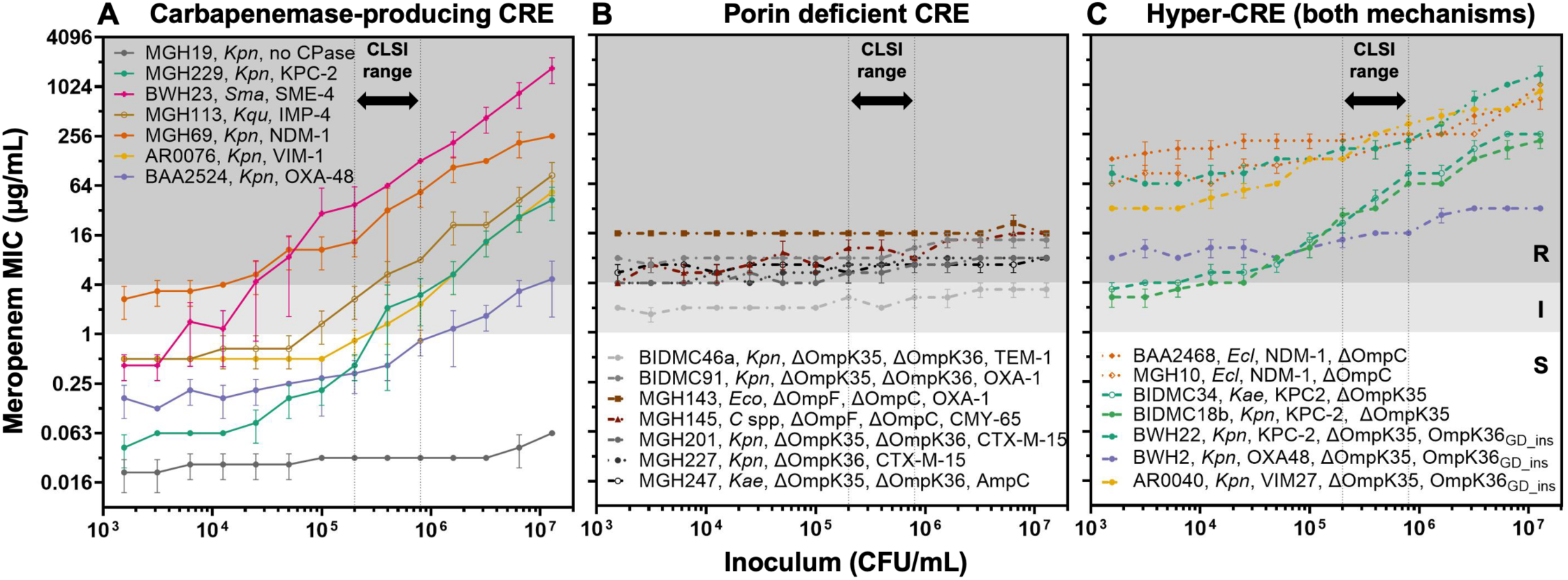
Select CRE exhibit different patterns of carbapenem inoculum effect depending on resistance mechanism. Meropenem MICs from broth microdilution assays (y-axis) across a broad range of starting inocula (x-axis) of select clinical CRE isolates encoding **(A)** a carbapenemase of each major type, **(B)** porin deficiency, or **(C)** both, with species and relevant genotype as indicated. One susceptible isolate (MGH19, gray) is also shown in panel **(A)**. Each MIC point is the mean of three replicates (error bars = standard error of the mean). *C spp* = *Citrobacter* spp, *Ecl* = *Enterobacter cloacae*, *Eco* = *Escherichia coli*, *Kae* = *Klebsiella aerogenes*, *Kpn* = *Klebsiella pneumoniae*, *Kqu* = *Klebsiella quaispneumoniae*, *Sma* = *Serratia marcescens*. CPase = carbapenemase. Vertical dotted lines reflect the CLSI-recommended inoculum range, background shading reflects CLSI breakpoints, and line color reflects carbapenemase content, all as in Fig. 1; dashed lines indicate porin deficiency.

### Carbapenemase production and porin deficiency together cause a hyper-CRE phenotype

To directly test the hypothesis that the combination of carbapenemase production and porin deficiency confers a hyper-CRE phenotype, we selected two clinical *Klebsiella pneumoniae* isolates (BIDMC46a and BIDMC91) with disruptions in both major porins OmpK36 and OmpK35, and transformed them with a pBAD33 plasmid encoding a KPC-3 (Fig. 3*A*). Both transformants exhibited dramatic increases in MIC at standard inoculum (256-fold and 128-fold, respectively) compared to empty vector controls and also developed an IE (6-fold from standard to high inoculum). In a second experiment, we disrupted the major OmpK36 porin in a *K. pneumoniae* isolate expressing a KPC-2 (RB582) using a CRISPR-Cas9 base editor (56). Meropenem MICs at standard inoculum increased by 32-fold in the edited strain, with a small IE (2-fold from standard to high inoculum, Fig. 3*B*). These experiments provide direct evidence that the combination of carbapenemase production and porin deficiency leads to a hyper-CRE phenotype.

**Fig. 3.**
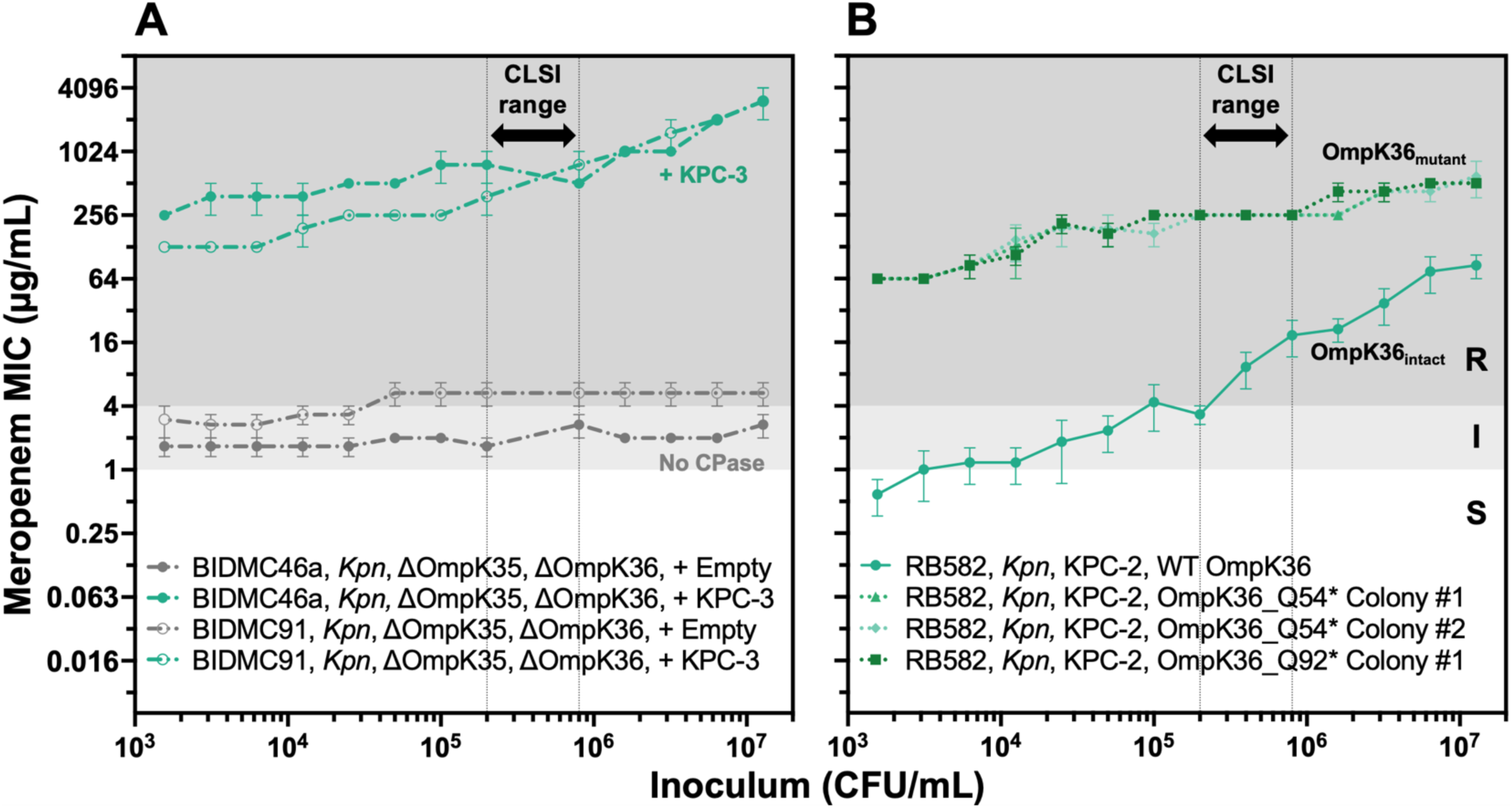
Carbapenemase production and porin deficiency together create a hyper-CRE phenotype. Broad-range meropenem broth microdilution assays for **(A)** two clinical PD-CRE isolates before and after transformation with pBAD33_KPC-3 and **(B)** a clinical CP-CRE isolate and three derivative colonies in which OmpK36 was disrupted by CRISPR-Cas9 based gene editing to create a premature stop at one of two glutamines as indicated. The parental KPC-expressing isolate (RB582) is shown as a solid line with green markers. Each MIC point is the mean of three replicates (error bars = standard error of the mean). Vertical dotted lines reflect the CLSI-recommended inoculum range, background shading reflects CLSI breakpoints, and line color reflects carbapenemase content as in Fig. 1; dashed lines indicate porin deficiency as in Fig. 2. *Kpn* = *Klebsiella pneumoniae*.

### Antibiotic exposure enhances CP-CRE release of carbapenemase activity into media

The strict dependence of the IE upon resistance genotype suggests that carbapenemase production may directly cause the inoculum effect. One plausible molecular mechanism is that carbapenemases are shared as common goods protecting bacterial populations (57). To test this hypothesis, we set up supernatant transfer experiments (Fig. 4*A*) to ask whether CP-CRE, or PD-CRE as controls (each referred to as “producer strains” in this experimental setup), release carbapenemase activity that can protect a fully susceptible reporter strain, *E. coli* K-12. Surprisingly, supernatant from cultures of three clinical CP-CRE grown to high inoculum (1 x 10^7^ CFU/mL), each from a different species and expressing a different carbapenemase, had no effect on reporter strain MIC at baseline (Fig 4*B* top panel, dashed lines). However, upon treatment with meropenem, supernatant from these high-inoculum cultures protected the K-12 reporter strain, increasing its meropenem MIC by up to >100-fold (Fig. 4*B* top panel, solid lines). By contrast, supernatant from high-inoculum PD-CRE cultures had no effect on the meropenem MIC of the reporter strain, with or without meropenem treatment (Fig. 4*B* bottom panel, solid vs dashed lines, respectively). Notably, we treated each producer strain with meropenem at half its respective standard-inoculum MIC, which for the tested CP-CRE strains was 16-to 128-fold below their MIC at the high inoculum at which we treated them, but 1-to 8-fold above their low-inoculum MIC (these relationships varied depending on the shape of the IE curve for each strain). Despite treating so far below each producer strain’s MIC at the treated (high) inoculum, we nonetheless observed death or reduced growth of CP-CRE strains in the initial 2 hours of treatment (Supplementary Fig. S3).

**Fig. 4.**
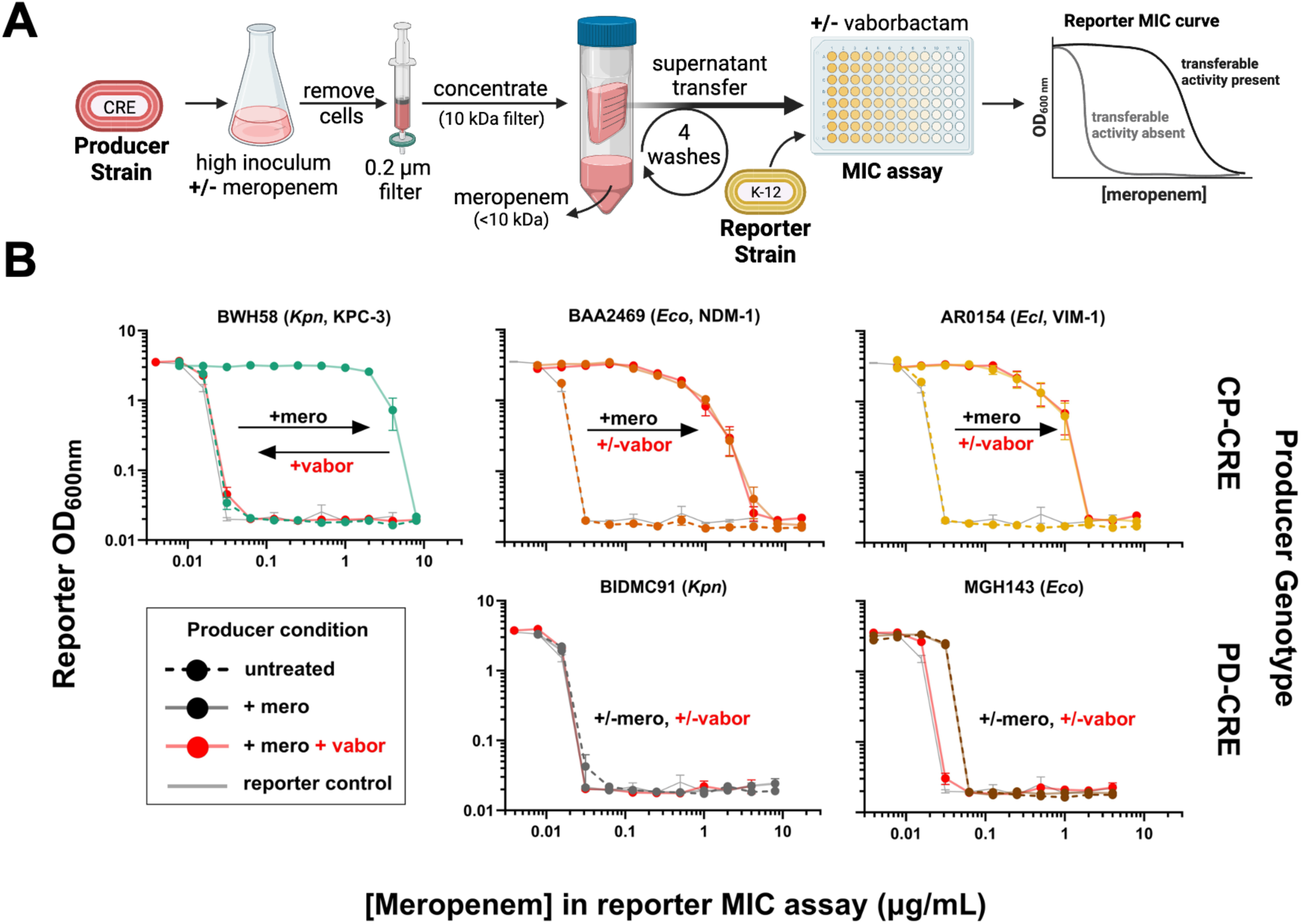
Clinical CP-CRE generate transferrable carbapenemase activity upon meropenem treatment. **(A)** Supernatant transfer experiment workflow: CP-CRE or PD-CRE isolates (“producer strains”) were grown to high inoculum, then exposed to media with or without antibiotic for 2 hours. Culture supernatants were filtered, washed to remove antibiotic, concentrated, and added to meropenem MIC assays for susceptible *E. coli* K-12 reporter strains to assess for transferrable protection. Mock data on the right shows expected results from reporter MIC assays if the supernatant contains protective activity (black) or not (gray). **(B)** Plots of reporter *E. coli* K-12 meropenem MIC assays, supplemented with supernatants from producer strains treated with (solid lines) or without (dashed lines) meropenem prior to supernatant transfer. CP-CRE producers are colored by carbapenemase content as in Fig. 1; gray curves = MIC of reporter strain alone; red curves = vaborbactam added to reporter MIC assay. Data from untreated samples and treated PD-CRE producers are the mean of three replicates of reporter MIC assays from a single producer treatment. All other data are the mean of nine replicates, three reporter MIC assays from each of three producer treatment replicates (error bars = standard error of the mean). *Ecl* = *Enterobacter cloacae*, *Eco* = *Escherichia coli*, *Kpn* = *Klebsiella pneumoniae;* mero = meropenem, vabor = vaborbactam. Isolates are colored by carbapenemase content as in Fig. 1.

To test whether this phenomenon generalized to all carbapenemases, we repeated these supernatant transfer experiments using our laboratory strains of *E. coli* K-12 transformed with each carbapenemase as producer strains (Supplementary Fig. S4). Each transformant generated transferrable carbapenemase activity after meropenem treatment (Supplementary Fig. S4*B*). In this laboratory strain background, unlike in the clinical producers, the stronger carbapenemases (all but OXA-48) generated detectable transferrable protection in the absence of antibiotics as well (dashed lines), though in all cases the amount of protection increased markedly with antibiotic treatment (solid lines).

This transferrable carbapenemase activity in the supernatant was released (from clinical CP-CRE strains) or enhanced (from K-12 transformants) not only by meropenem treatment, but also by treatment with the aminoglycoside apramycin (Supplementary Figs. S4*C* and S5), an unrelated antibiotic to which all strains were susceptible. Consistent with this protective activity coming from released carbapenemase, the selective carbapenemase inhibitor vaborbactam fully reversed the protection from KPC– and SME-producing strains, but not from strains producing the class B or D β-lactamases NDM, IMP, VIM, or OXA-48 (Fig. 4, Supplementary Figs. S4 and S5, red lines), matching vaborbactam’s known spectrum of activity (58).

### A significant proportion of carbapenemase-producing isolates change classification within the CLSI inoculum range

Although clinical microbiological laboratories initially identified the isolates we studied as carbapenem-resistant, the IE caused the MIC of several isolates to vary greatly within the CLSI inoculum range (Fig. 2*A*, and Supplementary Fig. S2*A*). This variability prompted us to quantify which carbapenemase-producing isolates changed susceptibility classification within the acceptable CLSI inoculum range of 2 × 10^5^ CFU/mL to 8 × 10^5^ CFU/mL. We tested both meropenem and ertapenem, two carbapenems in common clinical use with distinct physicochemical and pharmacokinetic properties. Notably, ertapenem has a lower barrier to resistance (59) since it is more affected by reduced outer membrane permeability (60), and has lower clinical susceptibility breakpoints (24). 22 of 63 (35%) carbapenemase-producing isolates tested with meropenem and 7 of 41 (17%) tested with ertapenem changed AST classification (Fig. 5). When excluding hyper-CRE and considering only isolates with intact porins, 18 of 36 (50%) CP-CRE isolates changed AST classification for meropenem and 7 of 29 (24%) for ertapenem within the CLSI inoculum range. Concerningly, three isolates changed from meropenem susceptible to resistant within the CLSI range (Fig. 4, dotted lines), including one *E. coli* isolate (BIDMC43b) and two *Serratia marcescens* isolates (BWH56 and BWH57), each of which encodes a KPC-3. Four isolates tested fully susceptible to meropenem throughout the CLSI inoculum range (red circle), including two *Citrobacter freundii* harboring KPC-3 (MGH281 and MGH283), and one *E. coli* (BAA2523) and one *K. pneumoniae* (BAA2524) each harboring an OXA-48. In all, 15 of 63 (24%) carbapenemase producers (CP-CRE or hyper-CRE) tested meropenem susceptible, and 2 of 41 (4.9%) tested ertapenem susceptible, at some point in the CLSI inoculum range; this represented 15 of 36 (42%) and 2 of 29 (6.9%) CP-CRE, respectively.

**Fig 5.**
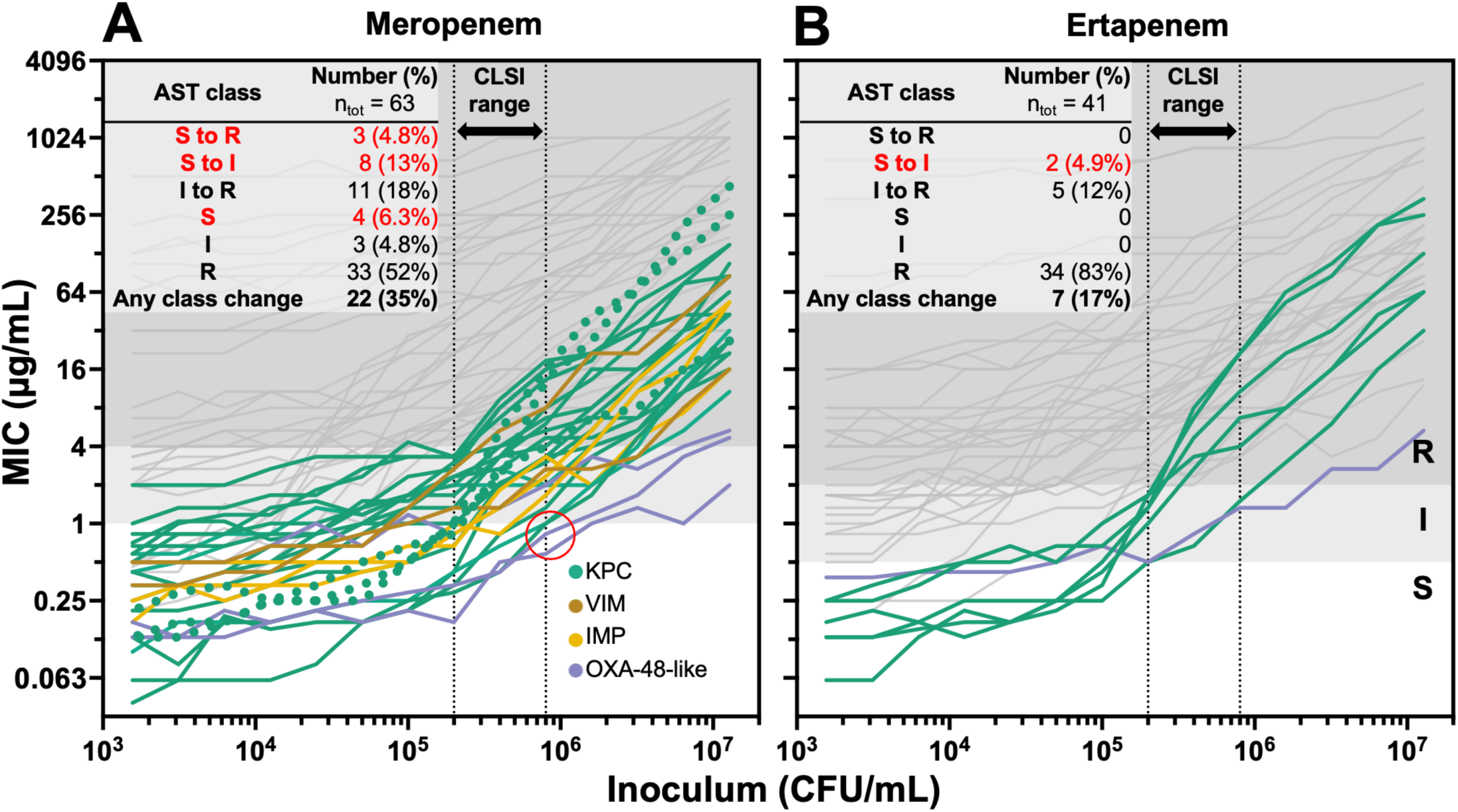
Many carbapenemase-producing isolates change AST classification in the CLSI range. Broad-range broth microdilution assays for **(A)** 63 carbapenemase-producing clinical CRE isolates (36 CP-CRE and 27 hyper-CRE) treated with meropenem and **(B)** 41 carbapenemase-producing clinical CRE isolates (30 CP-CRE and 11 hyper-CRE) treated with ertapenem. Each MIC measurement was the mean of three replicates; error bars were omitted for simplicity. Isolates that tested anything other than resistant at some point in the CLSI-recommended inoculum range are colored by carbapenemase content; those that tested resistant throughout the CLSI range are in gray. Inset table shows the number and percentage of isolates by AST class across the CLSI inoculum range; red text indicates isolates that tested susceptible at some point within the CLSI range. The red circle highlights 4 isolates that tested fully meropenem-susceptible throughout the CLSI range; dotted lines indicate 3 isolates whose meropenem susceptibility category changed from susceptible to resistant in the CLSI range.

### The inoculum effect accurately identifies carbapenemase production among CRE

The meropenem MIC at standard inoculum poorly distinguished carbapenemase-producing isolates (CP-CRE or hyper-CRE) from PD-CRE (Fig. 6*A*), making it an inefficient metric to trigger carbapenemase testing. By contrast, the clear phenotypic differences across inocula between these groups led us to investigate a diagnostic approach based on the IE. We quantified the IE by dividing the MIC at high inoculum (1.3 × 10^7^ CFU/mL) by the MIC at standard inoculum (4 × 10^5^ CFU/mL) for each isolate (Fig. 6*B*). CRE exhibited distinct meropenem IEs depending on their mechanism of resistance, with carbapenemase-producing isolates (CP-CRE and hyper-CRE) exhibiting a much more prominent IE, with a median increase in MIC of 10-fold, compared to a 1.2-fold increase for PD-CRE (Fig. 6*B*). To quantify binary classification distinguishing carbapenemase-producing isolates from porin-deficient ones, we generated receiver operating characteristic (ROC) curves using meropenem MIC (Fig. 6C) or IE (Fig. 6D) threshold values. The area under each ROC curve (AUC) represents an aggregate measure of classification accuracy in distinguishing isolates by resistance mechanism across all possible binary thresholds of the metric being plotted. The meropenem IE distinguished carbapenemase-producing CRE from PD-CRE better than meropenem MIC at standard inoculum, illustrated by a ROC AUC of 0.99 for the meropenem IE versus 0.69 for meropenem MIC. To efficiently identify CRE that might carry carbapenemases, precise IE threshold values can be chosen to optimize sensitivity (detecting carbapenemases) while maintaining reasonable specificity (red points in Fig. 6*D*). At one specific IE threshold of 2.53-fold, the sensitivity and specificity of distinguishing carbapenemase-encoding isolates from PD-CRE was 95.2% and 95.7%, respectively. The meropenem IE was greater in CP-CRE (median 16-fold MIC increase) than hyper-CRE (median 5.3-fold), and it also distinguished these two groups (Fig. 6*B* and 6*D*; ROC AUC 0.89).

**Fig. 6.**
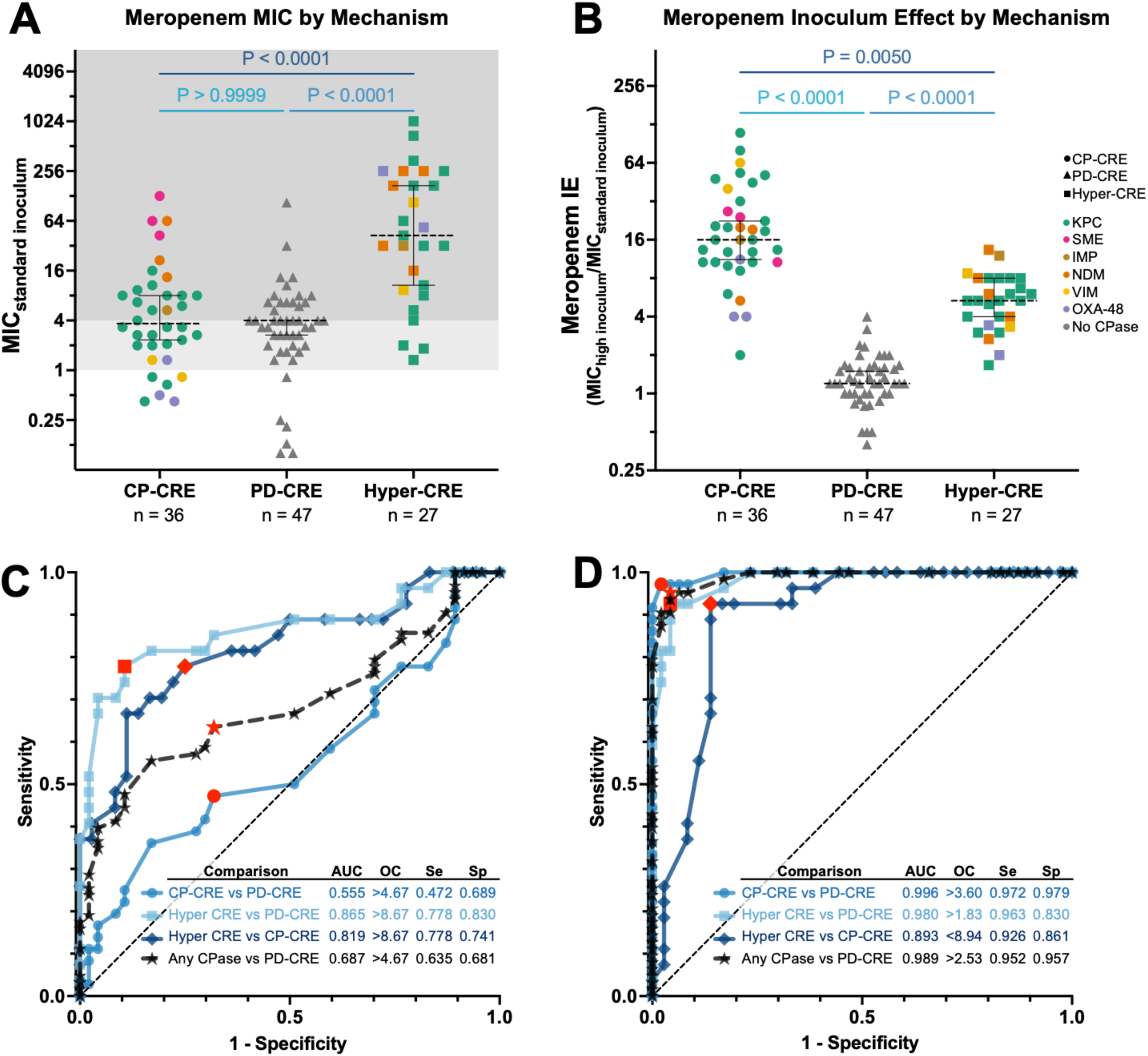
Meropenem inoculum effect robustly classifies CRE isolates by resistance mechanism. **(A)** Meropenem MICs for 106 CRE and 4 porin-deficient CSE (PD-CSE), grouped by genotypic mechanism of carbapenem resistance: CP-CRE (circles), PD-CRE or PD-CSE (triangles), or hyper-CRE (squares). Isolates are colored by carbapenemase content as in Fig. 1. Each MIC point is the mean of three replicates. **(B)** Ratio of meropenem MIC at high (1.3 x 10^7^ CFU/mL) versus standard (4.0 x 10^5^ CFU/mL) inoculum for 106 CRE and 4 PD-CSE, grouped as in **(A).** P-values were obtained by conducting statistical analyses on the three groups in **(A)** and **(B)** using a Kruskal-Wallis test. **(C)** Receiver operating characteristic (ROC) curves based on the meropenem MICs at standard inoculum from **(A)** comparing the different genotypic CRE groups. Inset tabulates each ROC area under the curve (AUC), sensitivity (Se), and specificity (Sp) at an optimal cutoff (OC) threshold, indicated by the red points on the ROC curves. **(D)** ROC curves based on the meropenem inoculum effect, displayed as in **(C)**. Any CPase = either CP-CRE or hyper-CRE.

### Comparing ertapenem and meropenem MICs gives insights into resistance mechanisms

The meropenem IE effectively distinguished carbapenemase-producing isolates from PD-CRE, but it is not a standard workflow in clinical laboratories, which typically only perform AST at standard inocula. To overcome this limitation, since the periplasmic permeability of ertapenem is typically more impacted by porin mutations than that of meropenem (6, 7, 61, 62), we hypothesized that comparing ertapenem and meropenem MICs, even at a single inoculum, might also distinguish CRE by mechanism of resistance. We thus measured ertapenem MICs across a broad inoculum range for 85 of the 110 CRE isolates (Fig. 7). The ratio of ertapenem MIC to meropenem MIC better separated carbapenemase producing isolates (CP-CRE or hyper-CRE) from PD-CRE (Fig. 7*C*) than either the ertapenem or meropenem MIC alone (Fig. 7 *A* and *B*). At standard inoculum, meropenem MIC performed poorly at distinguishing carbapenemase producers (CP-CRE or hyper-CRE) from PD-CRE (ROC AUC 0.559), whereas ertapenem MIC performed slightly better (ROC AUC 0.696; Fig. 7*D*). By contrast, the ertapenem-to-meropenem MIC ratio yielded a ROC AUC of 0.941, with a 90% sensitivity and 84% specificity at a ratio cutoff of 7.63 for detecting carbapenemase content. At low inoculum, ertapenem MIC performed comparably to this ratio at identifying carbapenemase producing isolates, while at high inoculum, meropenem MIC performed comparably. However, at the standard CLSI inoculum, neither performed as well as the MIC ratio.

**Fig. 7.**
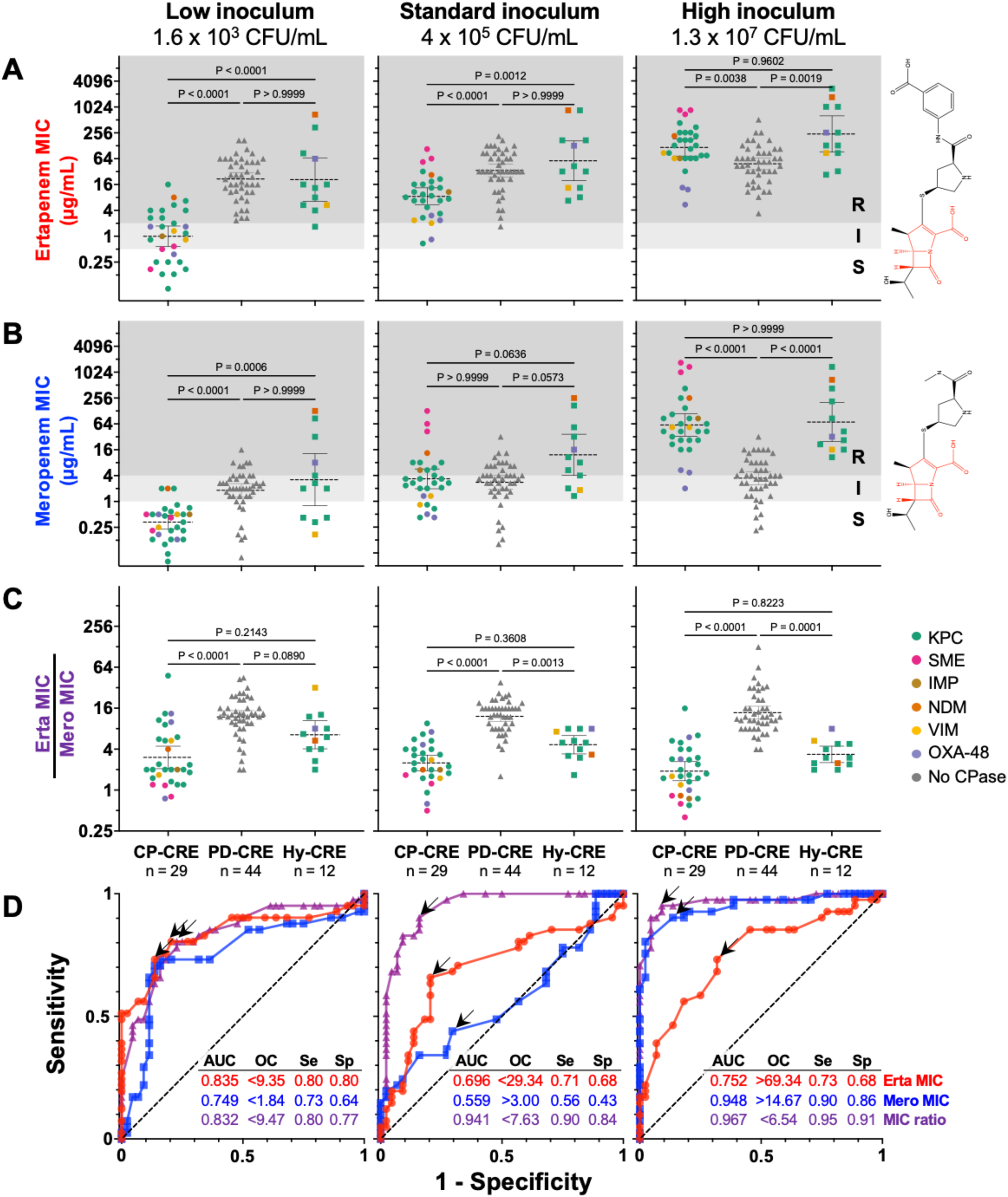
Comparing ertapenem and meropenem MICs gives insight into resistance mechanism. (**A-C**) MICs of ertapenem **(A)** or meropenem **(B)**, or the ratio of the ertapenem MIC to the meropenem MIC **(C),** are shown for 85 isolates at low, standard, and high inoculum (left, middle, and right panels respectively), grouped by resistance mechanism along the x-axis (hyper-CRE = “Hy-CRE”). Data points are colored based on carbapenemase content as in Fig. 1. Plot backgrounds are shaded by CLSI breakpoints for ertapenem **(A**) or meropenem **(B**) as in Fig. 1. P-values reflect non-parametric Kruskal-Wallis tests for each between-group comparison. In chemical structures, the carbapenem core is red and the modifiable substituents are black. **(D)** ROC curves comparing carbapenemase producers (CP-CRE + hyper-CRE) vs PD-CRE for each metric (red = ertapenem MIC, blue = meropenem MIC, purple = MIC ratio) at low, standard, and high inoculum. Inset tabulates ROC AUC and sensitivity (Se) and specificity (Sp) at an optimal cutoff (OC, indicated by arrow on ROC curve).

## Discussion

Our systematic investigations revealed that the *in vitro* carbapenem inoculum effect (IE) is not determined by the mechanism of action of the antibiotic, but rather by the mechanism of resistance of the isolate. In *E. coli* K-12, we showed that expression of any carbapenemase is sufficient to produce a carbapenem IE (Fig. 1). We confirmed this result in a collection of 106 clinical CRE isolates, in which all CP-CRE exhibited a clear IE, whereas PD-CRE did not (Fig. 2, Supplementary Fig. S2, and Table S3). Isolates with both mechanisms of resistance (hyper-CRE) tended to have higher MICs and also exhibited an IE, although smaller in magnitude than CP-CRE (Fig. 2*C* and Fig. 6*B*). Although a link between β-lactamase expression and the β-lactam IE has been long suspected, the link has not previously been established this clearly: in *S. aureus*, for instance, the mere presence of a β-lactamase is insufficient to predict a cefazolin IE (43–45). Indeed, *S. aureus* strains encoding the same family of *blaZ* β-lactamase may or may not exhibit a cefazolin IE, without obvious explanation after considerable study (43, 44), despite the absence of a periplasm that simplifies the system (63). Here, aided by the relatively limited and well-defined subset of β-lactamases that hydrolyze carbapenems, we were able to establish an obvious dichotomy in which carbapenemase content predicts a meropenem IE in the Enterobacterales.

These findings have important mechanistic, clinical, and diagnostic implications. Mechanistically, all carbapenemases imparted a clear inoculum effect despite limited sequence or structural similarity across the four Ambler classes. Furthermore, we observed a consistently biphasic dependence of CP-CRE MIC on inoculum (Fig. 1, Fig. 2 and Supplementary Fig. S2): at the lowest inocula tested, carbapenemases conferred a relatively constant degree of resistance, whereas above a certain threshold, resistance of the population increased with cell density, by up to 100-fold at the highest cell densities compared with the standard inoculum (Fig. 1B, Fig. 2A). This is consistent with a model where carbapenemases provide cell-autonomous resistance at low density but increasing population-level resistance as cell density increases, such that the collective production of carbapenemases may become the dominant determinant of resistance at high enough cell density. Since the IE is only observed with carbapenemase producers and not PD-CRE, one intriguing possibility is that carbapenemases serve as a “common good” in bacterial populations above a certain density threshold, as has been proposed for other secreted enzymes, including other β-lactamases (37–39, 64). One recent study of bacterial communities in which some members produced OXA-48 supports this model (40).

Our supernatant transfer experiments (Fig. 4, Supplementary Figs. S4 and S5) support this “common good” model but add complexity: carbapenemase release into the media is greatly enhanced upon antibiotic treatment, functionally consistent with antibiotic-mediated altruistic cell death, a phenomenon often discussed (65) but seldom confirmed outside of artificially engineered systems (66). This finding is consistent with a recent preprint proposing a model whereby AmpC-mediated cefotaxime hydrolysis may accelerate after cell lysis (67). Further studies are needed to more completely characterize this mechanism, including whether carbapenemase activity is released as free enzyme or enclosed in outer membrane vesicles (68–70), and whether or how its release is regulated upon antibiotic exposure, versus passively spilled by cell death. The enhanced release of carbapenemase activity by apramycin (Supplementary Fig. S5), an unrelated antibiotic class that inhibits ribosome function, implies that new protein synthesis is not required for release, perhaps favoring a passive release mechanism. Exploration of the implications of this shared activity for emergent resistance in CP-CRE populations compared with individual cells, both in laboratory culture and in structured and mixed bacterial communities that better reflect natural environments, is also warranted.

Notably, PD-CRE isolates expressing extended spectrum β-lactamases (ESBLs), which often contribute to carbapenem resistance, did not exhibit an appreciable carbapenem IE (Fig. 2*B* and Supplementary Fig. S2*B*), nor did several meropenem-susceptible isolates, including some encoding an ESBL and other randomly selected non-ESBL-producers (Supplementary Fig. S2*D*). Consistent with these observations, transformation of the common ESBL gene *bla*_CTX-M-15_ did not confer a meropenem IE upon *E. coli* K-12 (Supplementary Fig. S1). Together, these findings indicate that the slow rates of carbapenem hydrolysis by ESBL enzymes were insufficient to benefit neighboring bacteria even at high culture density. The lower prevalence of carbapenemases compared to other β-lactamases may explain why the IE is reported less frequently for carbapenems than for other β-lactams (31), but this might change as carbapenemases continue to spread globally.

The clinical implications of the inoculum effect have been uncertain because the relevant bacterial cell densities during most infections *in vivo* are generally below those used for AST (61, 64). For instance, in bacteremia, the infectious burden of ≤10 CFU/mL is several orders of magnitude below that at which the inoculum begins to affect MIC in our assays (71, 72).

Nevertheless, in *Staphylococcus aureus* bacteremia, a cefazolin IE has been associated with treatment failure and higher mortality (73, 74). Our findings suggest that at the low inoculum present in bacteremia, PD-CRE (Fig. 2*B*, Supplementary Fig. S2*B*) exhibit higher carbapenem MICs than CP-CRE (Fig. 2*A*, Supplementary Fig. S2*A*), yet clinical outcomes are similar or worse for CP-CRE bacteremia (75, 76). It is possible that the increased resistance of PD-CRE at physiologically relevant bacterial densities may be offset by fitness costs associated with reduced permeability. Further research is needed to fully understand the relationship between MIC, IE, resistance mechanisms, and clinical outcomes. Intriguing recent studies found that isolates encoding carbapenemases and displaying a large IE require higher meropenem concentration to prevent emergence of highly resistant mutants (39, 59). Since we observed the largest IE among CP-CRE with intact porins, perhaps these extremely resistant escape mutants reflect *de novo* porin mutations, creating hyper-CRE.

Diagnostically, accurate recognition of carbapenem resistance is crucial for infection control strategies and patient care. Our results confirmed prior reports that minor differences in inoculum can lead to categorical changes in susceptibility interpretation (48), particularly impacting the detection of CP-CRE, the isolates that carry the greatest implications for infection control. Even though the isolates we examined were selected because they tested as CRE, many changed classification within the CLSI-acceptable inoculum range and even appeared susceptible (Fig. 5), suggesting that clinical algorithms may miss some isolates encoding carbapenemases (77). In fact, meropenem and ertapenem MICs of all CP-CRE isolates varied significantly in this range. Our observations in *E. coli* K-12 (Fig. 1) reveal that strains harboring OXA-48, CMY-10, or even KPC might only test resistant if they possess other factors, such as reduced antibiotic permeability. The clinical significance of isolates that express a carbapenemase but test susceptible in the standard inoculum range is uncertain and requires further study. Nonetheless, caution is generally advised when using carbapenems in infections with carbapenemase-producers, even if they test susceptible, due to the lack of evidence supporting clinical outcomes in these cases (24, 78). Our findings in a set of isolates selected for meropenem nonsusceptibility suggest that other carbapenemase producers may have been missed by standard diagnostic workflows, which could contribute to poor patient outcomes and the global spread of carbapenem resistance.

The strong correlation between the carbapenem IE and genotype (Fig. 2) allows for improved predictions about resistance mechanisms from phenotypic testing. Although MICs were not designed to identify resistance mechanisms, carbapenem MICs are routinely used to prioritize isolates for molecular carbapenemase testing (79), though different MIC thresholds for testing are recommended in the US (24) and Europe (27). Our study revealed that the meropenem IE is a more sensitive and specific indicator of carbapenemase production in CRE than MIC at any single inoculum (Fig. 6). Although measuring the meropenem IE would require additional testing workflows, which are likely impractical in overworked clinical microbiology laboratories, we also found that integrating ertapenem and meropenem MICs from routinely generated AST data at standard inoculum can inform the likelihood of carbapenemase production (Fig. 6). The ratio of ertapenem to meropenem MIC predicts resistance mechanism better than either MIC alone, perhaps because the periplasmic permeability of ertapenem is more strongly affected by porin disruption than that of meropenem (6, 61, 62), while both are hydrolyzed at similar rates by carbapenemases (80, 81). Specifically, CRE with a high ratio of ertapenem to meropenem MIC (more resistant to ertapenem) are likely to be PD-CRE, whereas those with a low MIC ratio (similar resistance to both) are likely to have wild-type porins, and thus likely require a carbapenemase to achieve carbapenem resistance. As with the IE, hyper-CRE appear to fall in between, and tend to have higher MICs at any inoculum. Since the IE impacts both ertapenem and meropenem MICs similarly (Fig. 5), this ratio predicts resistance mechanism independent of inoculum, an advantage over either ertapenem or meropenem MICs alone (Fig. 7).

We propose that this ertapenem to meropenem MIC ratio might therefore be better suited than either MIC alone to prioritize isolates for molecular or enzymatic carbapenemase detection (20, 82, 83). Recognizing that MIC thresholds for susceptibility may miss carbapenemase-producing isolates, EUCAST recommends a 16-fold lower MIC threshold for carbapenemase screening than their clinical susceptibility breakpoint (27). This lower threshold will lead to more sensitive carbapenemase detection, but at lower specificity, requiring far more testing. If validated in further studies at this lower MIC threshold, the ertapenem to meropenem MIC ratio may enable optimization of strain selection for molecular carbapenemase testing, increasing screening efficiency below the susceptibility breakpoint. By more efficiently identifying carbapenemase-producing isolates, our work could facilitate implementation of recent advances showing that molecular detection of carbapenemases can accelerate administration of targeted therapies, such as β-lactamase inhibitor combinations (84), and decrease mortality (22). Moreover, this phenotypic approach could detect strains producing rare carbapenemases, such as SME or CMY-10, that are not routinely included in testing panels, akin to the Carba NP (24, 85) and mCIM (21) methods. This finding, in addition to its therapeutic value, could serve as a surveillance tool for the emergence and evolution of carbapenemases (86). This work does not aim to replace well-validated clinical carbapenemase detection methods. Instead, it illustrates how biological phenomena can offer further insights into resistance mechanism in a clinical context, perhaps improving the efficiency of deployment of existing carbapenemase detection assays.

This study has several limitations and areas requiring further study. First, the clinical isolates analyzed were mostly chosen because they tested carbapenem resistant, limiting the ability to determine the frequency of CP-CRE isolates that may be missed by routine testing due to the IE. Further studies are needed to assess the performance of our proposed assays on isolates with MICs at or below the susceptible breakpoint. Second, automated AST instruments may not provide precise MIC measurements for ertapenem and meropenem outside a narrow range around the susceptibility breakpoints, requiring further investigation into the practical utility of these lower-resolution MIC estimates in screening for CP-CRE. Third, while we found carbapenemases sufficient to confer a carbapenem IE, other factors such as heteroresistance, metabolic factors, quorum sensing, and target distribution may also contribute in some isolates or circumstances. However, carbapenemases predictably cause a robust IE without requiring other factors. Our findings reveal a clear pattern: as the inoculum increases above a certain threshold, we observe a steady linear rise in MIC (Fig. 2 and Supplementary Fig. S2). This pattern is less consistent with heteroresistance, an alternative explanation for the IE whereby a rare resistant subpopulation emerges from larger inocula. With heteroresistance, one would expect a sudden step-like increase in MIC as inoculum rises, as opposed to the gradual MIC increase that we consistently observed. Lastly, this study did not explore the basis of the IE for other antibiotics, nor how other processes like antibiotic efflux may contribute to carbapenem resistance. However, the consistent relationship between resistance mechanism and the IE for carbapenems, where hydrolytic enzymes and other resistance mechanisms are relatively few and well characterized, may offer a useful general model for one major mechanism underlying the IE. Future studies could involve pairing these detailed phenotypic data and genotypic information (Supplementary Table S3) with precise enzymological parameters and direct quantification of enzymes and metabolites in fractionated cytoplasm, periplasm, vesicles, and supernatant in order to enhance and experimentally test existing quantitative models (41, 64). This would enable a more comprehensive description of how different mechanisms of antibiotic resistance impact the growth of bacterial populations when exposed to antibiotic selection.

Antibiotic resistance often reflects several molecular processes acting in concert. Carbapenemase content and porin status together explained the levels of carbapenem resistance in most isolates we tested. Some genera like *Klebsiella* appear to exhibit higher carbapenem MICs than others like *Citrobacter* even when they encode the same carbapenemase-containing plasmid (14). This may reflect baseline differences in permeability by phylogenetic grouping, which is consistent with general trends in the ertapenem-to-meropenem MIC ratio across species in our collection (Supplementary Table S3). However, some isolates were outliers, exhibiting unexpectedly high resistance or inoculum-dependent patterns that suggest uncharacterized elements contribute to their carbapenem resistance. In some cases, these unexpected behaviors were clues to simple errors in sample workup. For instance, in one isolate (MGH4), the deposited genome assembly did not reveal a carbapenemase (12) but it exhibited a strong meropenem MIC suggestive of a CP-CRE. We performed PCR and Sanger sequencing, revealing that MGH4 harbored a KPC, consistent with the IE. The plasmid was likely lost during strain passaging prior to whole genome sequencing. Detailed IE phenotype also allowed insights into porin polymorphisms of uncertain significance, which are a frequent challenge for genotypic prediction of β-lactam resistance (80, 82, 83, 87–91). Given the differential impact of porin mutations on each carbapenem, the ertapenem-to-meropenem MIC ratio may offer a phenotypic reflection of porin function. This simple parameter, derived from values already routinely measured, may thus be useful in assessing resistance mechanisms to carbapenems and other β-lactams. Even with the insight generated in this study, we observed that some of our CRE isolates had unexplained levels or patterns of carbapenem resistance across a broad range of inocula. Such outliers underscore remaining gaps in our knowledge and motivate further investigations to characterize unknown processes leading to high-level resistance, which might be encountered more frequently as CRE continue to spread globally.

## Methods

### Bacterial isolates

Most of the bacterial strains tested in this paper were collected from three major Boston-area hospitals (Beth Israel Deaconess Medical Center, BIDMC; Brigham and Women’s Hospital, BWH; Massachusetts General Hospital, MGH) and the University of California Irvine School of Medicine (UCI) as part of separate studies (12, 14), and approved by each hospital and the Broad Institute, adjudicated under Broad Office of Research Subject Protections (ORSP) designation ORSP-1833; additional IRB approval was granted by the Massachusetts Institute of Technology Committee on the Use of Humans as Experimental Subjects. Collection of additional isolates from the MGH and BWH clinical microbiology laboratories was approved by the Partners Health Care Institutional Review Board (IRB) under protocol 2015P002215. Additional isolates were acquired from existing publicly available collections through the Wadsworth Center of the New York State Department of Health (NYSDOH), the FDA-CDC Antimicrobial Resistance Isolate Bank (CDC, Atlanta, GA), and the American Type Culture Collection (ATCC) (Manassas, VA).

### Cloning of carbapenemases

*bla*_KPC-3_, *bla*_SME-2_, *bla*_IMP-4_, *bla*_NDM-1_, *bla*_VIM-27_, and *bla*_OXA-48_ along with their native promoters were cloned by Gibson assembly from isolates in our existing CRE collection (12, 14) encoding these carbapenemases into a pBAD33 plasmid which encodes chloramphenicol resistance (*SI Appendix*, Table S1). *bla*_CMY-10_ with its native promoter was synthesized in the same pBAD33 vector backbone by GeneWiz (Suzhou, China), since we did not have a clinical isolate expressing this enzyme.

### Broad-range broth microdilution

Our MIC assays were adapted from previously published broth microdilution methods (23, 92). See Supplemental Methods for details on setup of the broad inoculum range in this assay format.

### Supernatant transfer experiments

The producer strain (see Fig. 4*A* for schematic) was grown to logarithmic phase (OD_600_ 0.1 – 0.4) in MHB (Sigma, 70192), then back-diluted to a target inoculum of 1.3e x 10^7^ CFU/mL in 16 mL and incubated with or without antibiotic for 2 hours at 37°C on a shaking platform (250 rpm). Meropenem treatments were at 1/2x and apramycin treatments were at 4x the standard-inoculum MIC of each producer strain. After incubation, cultures were pelleted at 5000xg for 5 minutes at 4°C. Supernatants were syringe-filtered with 0.2 μm cellulose acetate filters (Cytiva, 6901-2502) to remove remaining producer cells, transferred to 10 kDa Amicon Ultra-15 centrifugal filter units (Millipore Sigma, UFC901025) pre-rinsed in phosphate-buffered saline (PBS), and spun at 5000xg at 4°C until concentrated to 1/10 of the original volume. Concentrated supernatants from meropenem-treated conditions were washed 3 times to remove meropenem (which would otherwise interfere with the reporter MIC assay) by adding PBS to the original volume and re-concentrating by 10x, then underwent a final MHB wash. This 10x-concentrated conditioned media was then added at a final 1x concentration to a standard meropenem broth microdilution MIC assay, with the reporter strain at standard inoculum (4 x 10^5^ CFU/mL) in MHB (Difco, 275730). For experiments with meropenem-treated producer strains (Fig. 4, Supplementary Fig. S3*A*), the reporter strain was *E. coli* K-12 MG1655; for apramycin experiments (Supplementary Figs. S3*B* and S4), the reporter strain was *E. coli* K-12 MG1655 transformed with an empty vector marked with an apramycin resistance gene (pBAD33-AprR), to eliminate the need for wash steps to remove the antibiotic from the producer supernatant. Vaborbactam (Cayman Chemical, 23962) was added at a final concentration of 8 μg/mL in select reporter MIC assays. See Supplemental Methods for further details.

### Data analysis

MIC replicates were imported to GraphPad Prism v 9.5.1 (GraphPad Software Inc., San Diego, CA); MIC vs inoculum plots (Figs. 1, 2, and 3, and Supplementary Fig. S1) depict mean and standard errors of the mean. To analyze the meropenem IE, the ratio of mean MIC at 1.3 × 10^7^ CFU/mL to 4 × 10^5^ CFU/mL was calculated. The IE of the 47 porin deficient CRE strains were compared to the 63 carbapenemase-encoding CRE strains using a non-parametric test (Mann-Whitney for comparisons between two groups or Kruskal-Wallis for analysis of more than two groups) in GraphPad Prism. Statistical analysis comparing PD-CRE isolates versus CP-CRE and hyper-CRE isolate at low, standard, and high inocula for ertapenem, meropenem, and ertapenem-to-meropenem MIC ratio was conducted using Mann-Whitney tests. All receiver operating characteristic (ROC) curves and analysis were constructed using GraphPad Prism.

### Gene editing to disrupt OmpK36 in *Klebsiella pneumoniae* isolate RB582

A CRISPR-Cas9 cytidine base-editing system (56) was used to introduce an early termination codon in the intact *ompK36* of isolate RB582. See Supplemental Methods for further details.

## Author Contributions

Designed research: AJC, KLT, LCL, JS, RPB

Performed research: AJC, KLT, LCL, JS, JDC, RPB

Contributed new reagents/analytic tools: AJC, KLT, AME, RPB

Analyzed data: AJC, KLT, LCL, JS, JTS, ALM, VMP, AME, RPB

Wrote the paper: AJC, KLT, LCL, JS, JTS, ALM, VMP, AME, RPB

## Classification

Biological Sciences / Microbiology

## Acknowledgements

We thank Pierre O. Ankomah, Nicoletta Commins, Alyssa DuBois, Alasdair Fletcher, James E. Gomez, Sarah Gomez-Villegas, Ria Kolli, Jonathan Livny, Peijun Ma, Melanie A. Martinsen, Michelle E. Matzko, David J. Roach, and Eleanor L. Young for helpful input.

We thank Kimberly Musser, the Wadsworth Center, and the CDC ARBank for sharing isolates.

This work was supported in part by the Broad Institute Next Generation Fund (to RPB), the Defeating Antibiotic Resistance through Transformative Solutions (DARTS) award from the Advanced Research Projects Agency for Health (to RPB), and federal funds from the National Institute of Allergy and Infectious Diseases, National Institutes of Health, Department of Health and Human Services, under Grant Number U19AI110818 to the Broad Institute (to AME). The content is solely the responsibility of the authors and does not necessarily represent the official views of any funding agency.

