## Supplementary Information for "The carbapenem inoculum effect provides insight into the molecular mechanisms underlying carbapenem resistance in Enterobacterales"

This file includes:
Supplementary Materials and Methods
Supplementary Figures S1-S5
Supplementary Tables S1 and S2
Legend for Supplementary Table S3 (.xlsx file)
Supplementary References

### Materials and Methods

**Genotypic determinants of carbapenem resistance in bacterial isolates.** Antibiotic resistance genes and porin status of BIDMC, BWH, MGH and UCI isolates were determined in previous work and are publicly accessible from https://www.ncbi.nlm.nih.gov/pathogens/ (1, 2). In brief, this previous work combined data from Illumina fragment libraries with either long-insert mate-pair Illumina sequencing libraries or Oxford Nanopore long-read libraries (2) to assemble high quality reference genomes, which were queried against multiple antimicrobial resistance databases. For porin status, the previous work identified porin sequences by similarity search against reference *ompC* and *ompF* sequences. These were classified as intact if they produced a full-length protein or as disrupted if they had frameshifts affecting 30 codons, truncations, transposon insertions in CDS or promoter, complete deletion (missing from WGS), or < 90% of the reference sequence. Additionally, *ompK36* was considered disrupted if it included codons encoding a glycine-aspartate insertion (3) or a C25T nucleotide mutation (4) that were previously shown to increase carbapenem resistance. Antibiotic resistance genes for FDA-CDC and ATCC are also publicly available from https://wwwn.cdc.gov/arisolatebank/ and https://genomes.atcc.org/. For FDA-CDC and ATCC isolates, publicly available genome assemblies were analyzed to determine if *ompC* (*ompK36*) and *ompF* (*ompK35*) or their respective homologs in other species were disrupted. For NYSDOH isolates, PCR was performed by NYSDOH for carbapenemase genes, and we performed Sanger sequencing to determine porin status.

**Gibson assembly cloning of carbapenemases**. $\textit{bla}_{\text{KPC-3}}$, $\textit{bla}_{\text{SME-2}}$, $\textit{bla}_{\text{IMP-4}}$, $\textit{bla}_{\text{NDM-1}}$, $\textit{bla}_{\text{VIM-27}}$ (5), and $\textit{bla}_{\text{OXA-48}}$ were cloned from isolates encoding these carbapenemases into a pBAD33 plasmid which encodes chloramphenicol resistance (Table S1). In brief, PCR primers were designed to amplify the carbapenemase genes to include their native promoters and terminators including 5' and 3’ overlapped ends for Gibson assembly cloning into a pBAD33 (6). The pBAD33 vector was also amplified to include 5’ and 3’ overlapped ends. The PCR reactions were digested by incubating with 1 µL of DpnI (NEB, MA, United States) at 37°C for 1 hour. The reaction products were purified using Zymo Research DNA Clean & Concentrator-25 (Zymo Research, Redwood Shores, CA). The concentration of the DNA products was measured using a NanoDrop ND-1000 spectrophotometer (Thermo Fisher Scientific, MA, United States) and assembled using a New England Biolabs (NEB) Gibson Assembly® Cloning Kit (Ipswich, MA). Next, 1 µL of the Gibson assembly reaction products were transformed into 50 µL NEB 5-alpha competent *E. coli* by incubating in ice for 15 minutes and then heat shocking the cells in a 42°C water bath for 30 seconds. The transformed cells were transferred to a 14 mL falcon tube with 0.5 mL of SOC media and the cells shaken at 200 rpm for 1 hour to allow for recovery before plating into LB agar plates supplemented with 100 µg/mL ampicillin. After overnight incubation, colonies from the plates were selected and the presence of the carbapenemases was verified using colony PCR. Plasmids were purified using a ZymoPURE Plasmid Miniprep Kit (Zymo Research, Redwood Shores, CA). The final plasmid constructs with the carbapenemases were verified using universal pBAD primers. Sanger sequencing revealed the pBAD33_KPC-3 construct had a K272H mutation, but it imparted levels of meropenem resistance on-par with those observed in other isolates encoding $bla_{\text{KPC-3}}$. The pBAD33_CMY-10 (7) was synthesized by GeneWiz (Suzhou, China). For MIC assays, the plasmids were purified and transformed into *E. coli* K-12 MG1655.

**Broad-range broth microdilution.** Our MIC assays were adopted from previously published methods (8). LB-agar plates supplemented with 100 µg/mL of ampicillin were streaked with a frozen glycerol stock of an isolate of interest to yield single colonies. After overnight growth, a single colony from the plate was selected and used to inoculate 2 mL of Mueller-Hinton Broth media (Becton, Dickinson and Company, Sparks, MD). The cultures were grown to exponential growth phase with an $OD_{600nm}$ of 0.3-0.8 The cultures were then diluted to the highest inoculum tested, estimating 1 OD to contain 8 × $10^8$ CFU/mL. The cultures were serially diluted by two-fold to yield the 14 inocula used from 1.3 × $10^7$ to 1.6 × $10^3$ CFU/mL. Meropenem and ertapenem (Combi-Blocks, San Diego, CA) were dissolved in MHB. MIC assays were conducted in 96-well plates where the antibiotics were serially diluted to then obtain the desired range of concentrations to capture the MIC for each strain at the 14 inocula yielding 100 µL of sample of each well. Each inoculum was also grown in the absence of antibiotics for subsequent normalization. After 16-20 hours in an ambient air incubator at 37°C, the absorbance of each well was read using a spectrophotometer, the OD was corrected to account for background signal, and MICs were determined as the lowest concentration that yielded ≥90% growth inhibition compared to the sample grown in the absence of antibiotics. We conducted our MIC experiments in triplicate with cultures inoculated with different colonies to account for the variability that can be present in broth microdilution (9), and all data reflect the average of each set of triplicates.

**Gene editing to knockout OmpK36 in *Klebsiella pneumoniae* isolate RB582.** A CRISPR-Cas9 cytidine base-editing system (10) was used to introduce an early termination codon in the intact *ompK36* of isolate RB582. The cytidine base-editing system (pBECKP_Apr plasmid) for precise *Klebsiella pneumoniae* C→T editing was ordered from Addgene (Watertown, MA). The protocol designed by Wang and colleagues was followed to generate the desired mutants: the pBECKP_Apr plasmid was linearized with an NEB BsaI HF (Ipswich, MA) and the product was cleaned up using a Zymo Research DNA Clean & Concentrator-25. Two different spacers serving as the CRISPR-Cas9 sgRNA template were designed to target two codons encoding glutamine amino acids (Q54 and Q92) suitably adjacent to PAM sites that could be recognized by a CRISPR-Cas9 system; 24 bp forward and reverse oligos were ordered from Azenta (Burlington, MA), phosphorylated with a T4 polynucleotide kinase (NEB), and ligated into the linearized pBECKP_Apr vector using a T4 DNA ligase (NEB). The ligation product was transformed into NEB 5-alpha competent *E. coli* competent cells as described above for the cloning procedures. After recovery, the transformed cells were plated into LB-agar plates supplemented with 50 µg/mL apramycin. The plasmid in the resulting colonies were verified with PCR and Sanger sequencing to contain the desired spacer. These colonies were used to inoculate cultures to purify the plasmids using a ZymoPURE Plasmid Miniprep Kit. This resulted in two plasmids to target the introduction of stop codons at two positions of OmpK36: $sgRNA_{Q54*}$ and $sgRNA_{Q92*}$.

To make RB582 competent cells, this strain was streaked onto an LB-agar plate supplemented with 100 µg/mL ampicillin. After overnight incubation, a single colony from the plate was used to inoculate 30 mL of LB broth. The culture was grown at 37°C and shaken at 250 rpm until the $OD_{600nm}$ reached 0.5. At that point the culture was pelleted and resuspended in deionized autoclaved water at 4°C. The cells were pelleted and resuspended two more times and the protocol described for transformation above was used to transform the $sgRNA_{Q54*}$ and $sgRNA_{Q92*}$ plasmids. After recovery, the transformed bacteria were plated into LB agar plates supplemented with 50 µg/mL apramycin. After overnight incubation, colonies were screened by PCR and Sanger sequencing to check if they had the desired mutations in *ompK36*. The $sgRNA_{Q54*}$ and $sgRNA_{Q92*}$ were cured by taking colonies with the desired mutations and inoculating them in LB broth. The cells were streaked into LB agar plates with 5% sucrose and grown.

**Transformation of pBAD33_KPC-3 into porin deficient isolates.** Competent cells for BIDMC46a and BIDMC91 were prepared as described above for RB582. Plasmid pBAD33_KPC-3 was purified and transformed into BIDMC46a and BIDMC91 as described in the cloning procedures. After recovery, the cultures were plated onto LB-agar plates supplemented with 34 μg/mL chloramphenicol. The presence of *bla*$_{KPC-3}$ in these strains was verified via PCR.

**Supernatant transfer experiment optimization:** In our initial experiments with the supernatant transfer assay (Fig 4*A*), we encountered media and filtration dependent variability in the observed results. Specifically, we first observed that the manufacturer of Mueller-Hinton Broth used to culture producer strains, but not reporter strains, impacted measured reporter MIC data. Culturing producer strains in Difco MHB (275730) resulted in no detectable transferrable protection with or without antibiotic treatment of the producer strain, whereas transferrable protection was reproducibly seen after antibiotic treatment when producer strains were cultured in Sigma MHB (70192). Varying the manufacturer of the media used in reporter MIC assays had no effect on results. To disentangle this effect from other steps in the assay, we varied multiple steps in the process and found that the activity was lost during 0.2 μm filtration of the producer strain supernatant prior to transfer to the reporter MIC assays. To isolate this step, we leveraged apramycin treatment experiments (as in Fig S5), using an apramycin-resistant reporter strain and performing reporter MICs in the presence of apramycin. This allowed us to omit the filtration step, which is otherwise required to prevent carryover of the meropenem-resistant producer strain, which could then overgrow in reporter MIC assays if not removed. Omitting the syringe filtration step restored transferrable carbapenemase activity from producer strains grown in Difco MHB, suggesting transferrable activity was initially present but lost during syringe filtration passages, rather than from lack of production, when producers were grown in Difco MHB. Minimal activity loss was observed from filtration of producers grown in Sigma MHB media, in contrast to Difco MHB media. Since the filtration step cannot be omitted in meropenem-treated producer strains, we used Sigma MHB for producer strain cultures in all supernatant transfer experiments for reproducibility.

**Supplementary Figures**

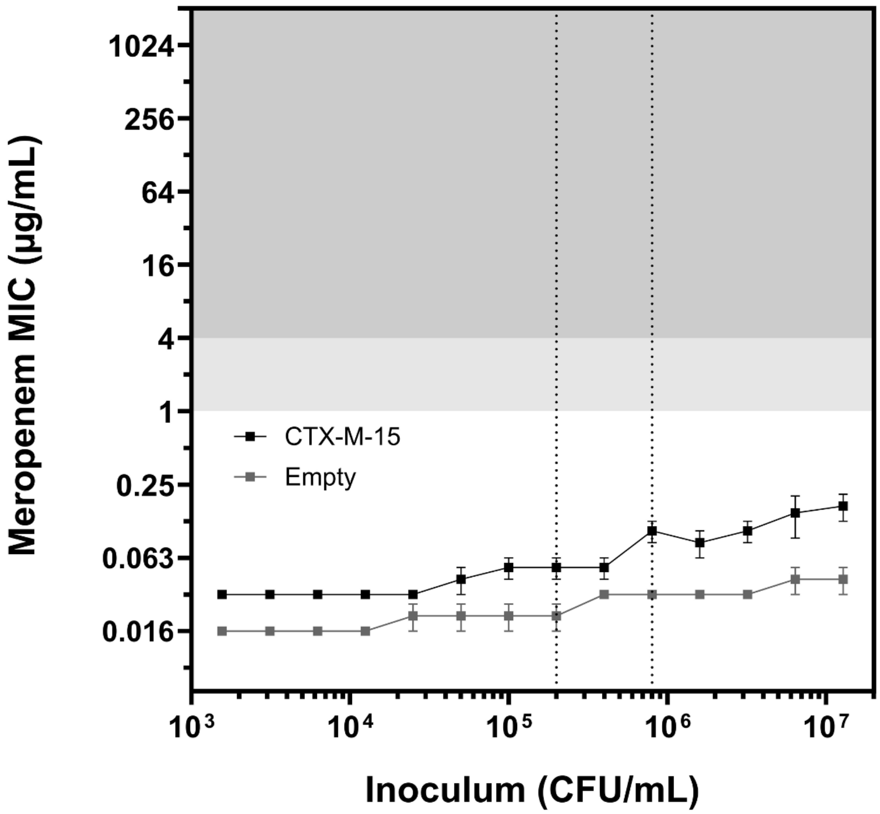


**Supplementary Fig. S1. Extended-spectrum β-lactamase CTX-M-15 imparts a minimal inoculum effect.** Broad-range meropenem broth microdilution for *E. coli* K-12 transformed with CTX-M-15 (black) or empty vector (gray). Each MIC point is the mean of three replicates (error bars = standard error of the mean). Vertical dotted lines represent the CLSI-recommended inoculum range (2 to 8 x $10^5$ CFU/mL). The background is shaded by CLSI meropenem susceptibility breakpoints (S = susceptible, white; I = intermediate, light gray; R = resistant, gray).

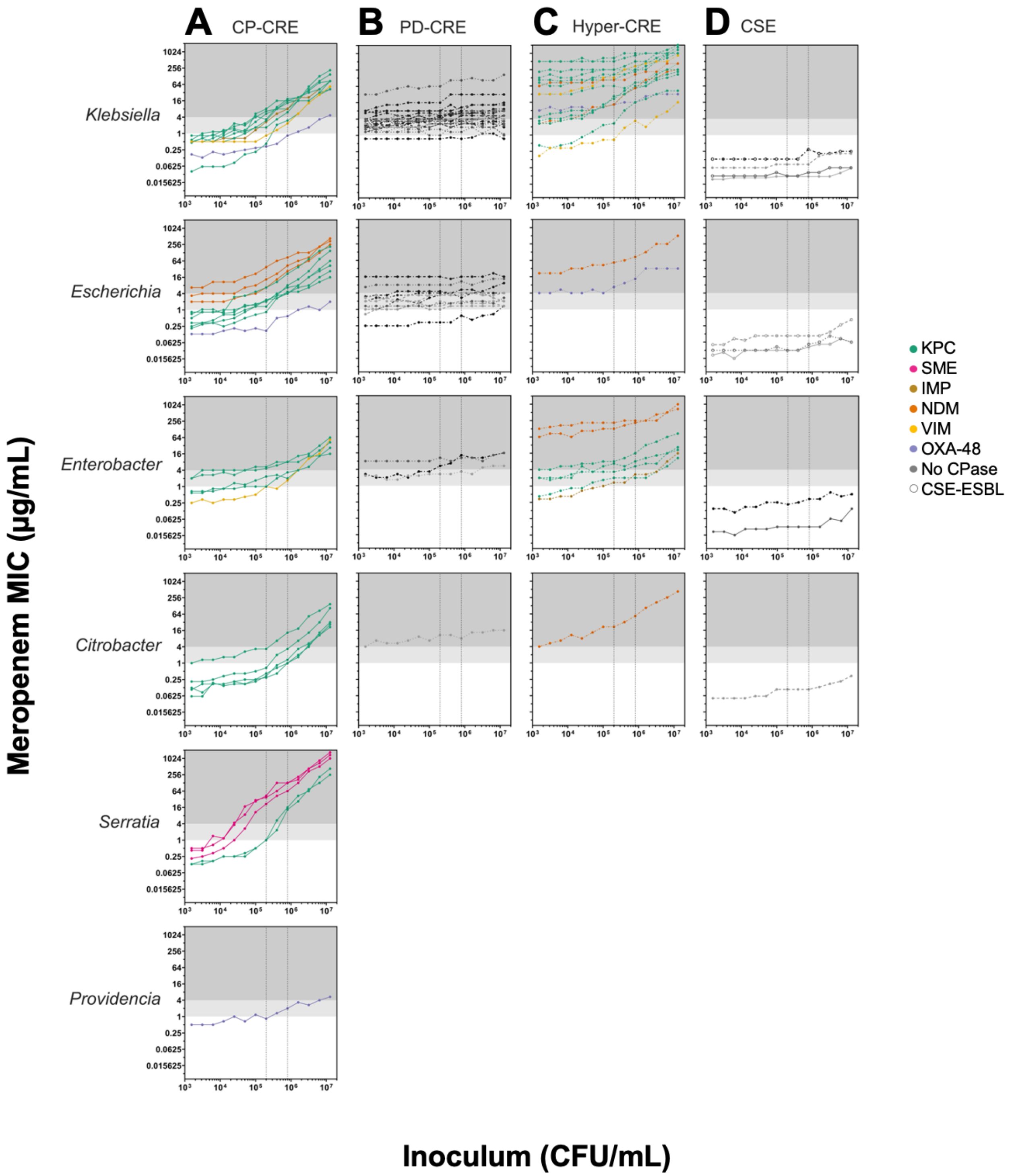


**Supplementary Fig. S2. MIC as a function of inoculum for 106 CRE of three resistance genotypes, and select CSE.** Broad-range meropenem broth microdilution MICs for **(A)** CP-CRE, **(B)** PD-CRE, **(C)** hyper-CRE, or **(D)** CSE separated vertically by genus. Each MIC point is the mean of three replicates; error bars were omitted for simplicity. Vertical dotted lines reflect the CLSI-recommended inoculum range, background shading reflects CLSI breakpoints, and line color reflects carbapenemase content as in Fig. 1; dashed lines indicate porin deficiency as in Fig. 2; open circles are ESBL-producing CSE (panel D only).

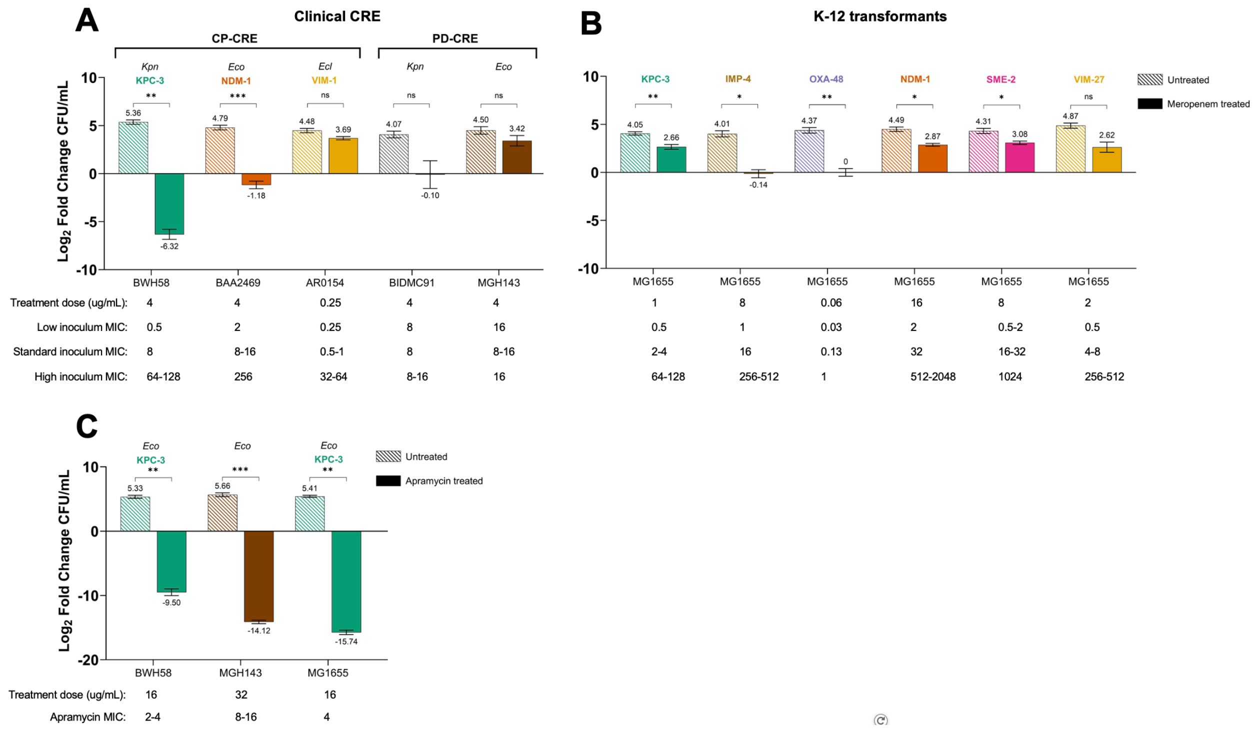


**Supplementary Fig. S3. Producer CRE populations are reduced by antibiotic treatment below the MIC at the treated inoculum.** Changes in cfu from producer cultures 2 hrs after incubation without (striped bars) or with (filled bars) antibiotic from supernatant transfer assays of meropenem-treated **(A)** clinical CRE (corresponding to Fig 4) or **(B)** K-12 transformant CP-CRE (corresponding to Fig S4), or **(C)** apramycin treated producer CRE (corresponding to Fig S5). Respective antibiotic treatments and measured MICs at low, standard, and high inoculum from Supplementary Table S3 tabulated underneath graphs for comparison. Data represent the mean of 1-3 replicates of producer strain treatment, as described in Fig 4, Fig S4, and Fig S5; error bars represent Poisson error from cfu counting, propagated through the computed standard deviation for producer strains with 3 treatment replicates. Statistical significance between untreated and treated conditions determined by Welch t-tests (ns = $p > 0.05$, * = $p < 0.05$, ** = $p < 0.01$, *** = $p < 0.001$). *Ecl* = *Enterobacter cloacae, Eco* = *Escherichia coli*, *Kpn* = *Klebsiella pneumoniae.* Isolates are colored by carbapenemase content as in Fig. 1.

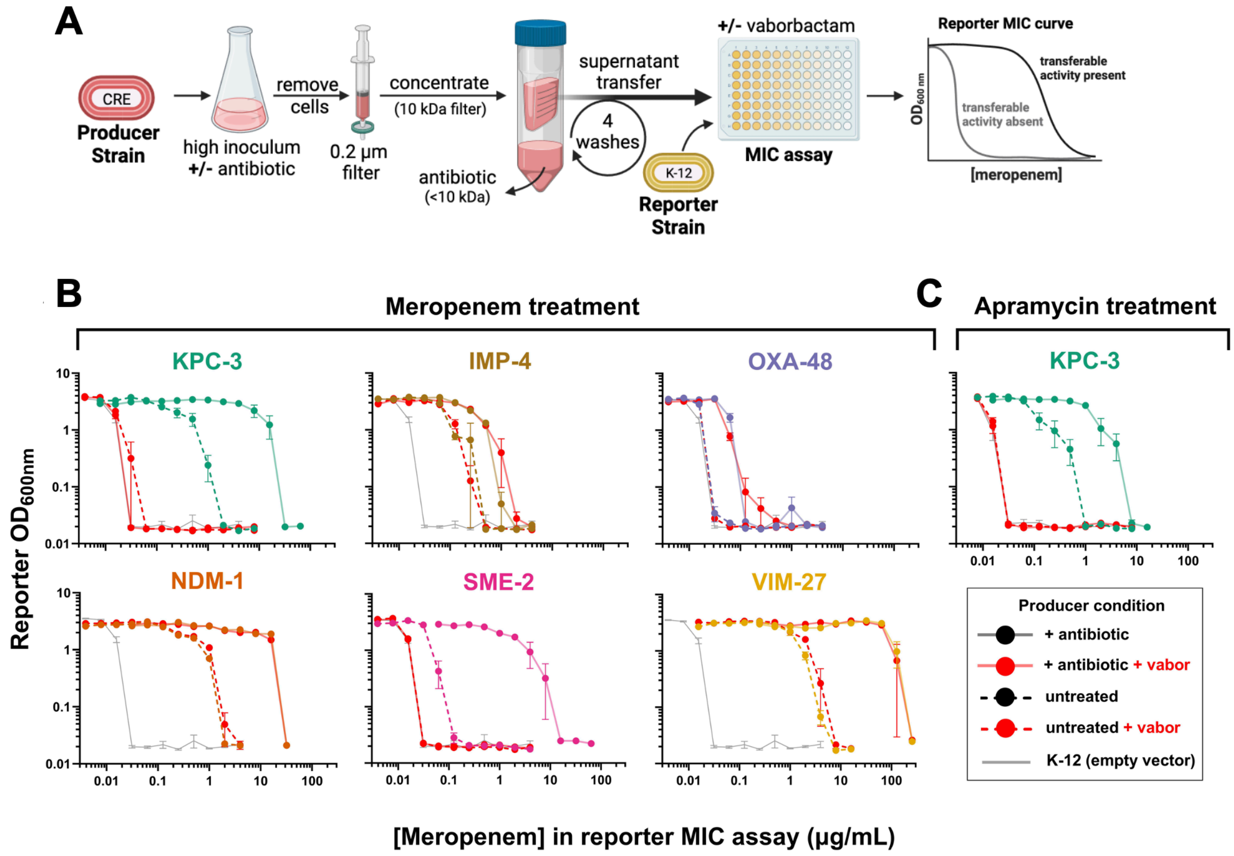


**Supplementary Fig. S4. Any carbapenemase class generates transferrable protection from *E. coli* K-12 upon antibiotic treatment.** Supernatant transfer experiments **(A)** were repeated as in Fig. 4, using *E. coli* K-12 transformed with the indicated carbapenemase as the producer strain. Producer cultures were treated (solid lines) with **(B)** meropenem or **(C)** apramycin or left untreated (dashed lines), and supernatants were transferred to an untransformed *E. coli* K-12 reporter strain for meropenem MIC assays as described in Fig. 4*A*. Curves are colored by producer carbapenemase content as in Fig. 1; gray curves = MIC of reporter strain alone; red curves = vaborbactam ("vabor") added to reporter MIC assay. Treated KPC-3 data are the mean of nine replicates, three reporter MIC assays from each of three producer condition replicates; all other data are the mean of three replicates of reporter MIC assays from a single producer condition (error bars = standard error of the mean).

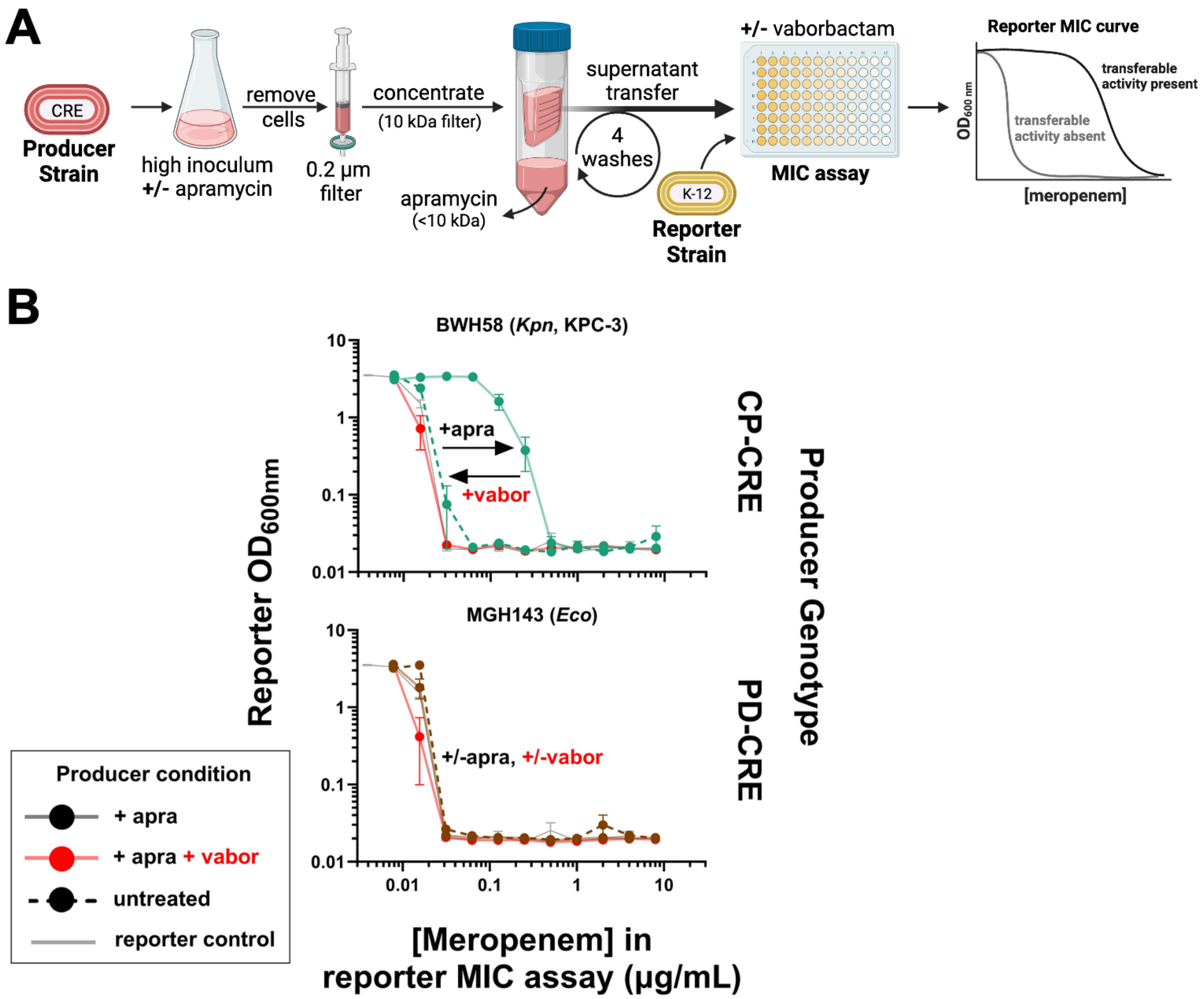


**Supplementary Fig. S5. Apramycin treatment of clinical CP-CRE exhibits transferrable carbapenemase activity to *E. coli* K-12. (A)** Supernatants were prepared and transferred to *E. coli* K-12 as in Fig. 4. **(B)** Meropenem MIC curve plots of reporter *E. coli* K-12 supplemented with supernatant harvested from producer strains under indicated apramycin conditions. Treated KPC-3 data are the mean of nine replicates, three reporter MIC assays from each of three producer treatment replicates; all other data are the mean of three replicates of reporter MIC assays from a single producer treatment (error bars = standard error of the mean). *Eco* = *Escherichia coli*, *Kpn* = *Klebsiella pneumoniae,* apra = apramycin, vabor = vaborbactam. Curves are colored by producer carbapenemase content as in Fig. 1.

### Supplementary Tables

**Table S1. Carbapenemases and sequences used for cloning**

| Carbapenemase cloned | Isolate template | Source reference | NCBI BioSample | Gene length | Upstream length | Downstream length | Total length |
|---|---|---|---|---|---|---|---|
| CMY-10 | YmcD1 | Lee et al. 2003 | SAMN12885507 | 1149 | 200 | 100 | 1449 |
| IMP-4 | MGH113 | Salamzade et al. 2022 | SAMN03280402 | 741 | 80 | 100 | 921 |
| KPC-3 | MGH283 | Salamzade et al. 2022 | SAMN08148336 | 882 | 406 | 137 | 1425 |
| NDM-1 | MGH69 | Salamzade et al. 2022 | SAMN02581242 | 813 | 104 | 100 | 1017 |
| OXA-48 | BWH2 | Salamzade et al. 2022 | SAMN02138589 | 798 | 163 | 250 | 1211 |
| SME-2 | BIDMC44 | Salamzade et al. 2022 | SAMN02356583 | 885 | 412 | 150 | 1447 |
| VIM-27 | AR0040 | Lutgring et al. 2018 | SAMN04014881 | 801 | 300 | 250 | 1351 |

**Table S2. Carbapenemase and porin status by genus of clinical isolates used in this study**

| | | Carbapenemases | | | | | | |
|---|---|---|---|---|---|---|---|---|
| Genus | Porin status | KPC | NDM | VIM | IMP | OXA48 | SME | None |
| *Citrobacter* | WT | 5 | - | - | - | - | - | 1 |
| | Deficient | - | 1 | - | - | - | - | 1 |
| *Enterobacter* | WT | 4 | - | 1 | - | - | - | 3 |
| | Deficient | 5* | 2 | - | 1* | - | - | 3 |
| *Escherichia* | WT | 6 | 3[†] | - | - | 2[†] | - | 1 |
| | Deficient | 1[‡] | 1[‡] | - | - | 1 | - | 14 |
| *Klebsiella* | WT | 7 | - | 1 | 1 | 1 | - | 2 |
| | Deficient | 12[§] | 3[§] | 2 | - | 1 | - | 29 |
| *Providencia* | WT | - | - | - | - | 1 | - | - |
| | Deficient | - | - | - | - | - | - | - |
| *Serratia* | WT | 2 | - | - | - | - | 3 | - |
| | Deficient | - | - | - | - | - | - | - |

*, †, ‡, § = an individual strain simultaneously carried the two indicated carbapenemases

**Table S3. Meropenem and ertapenem MIC metadata and genotype data for clinical isolates relevant for carbapenem resistance, including porin status, carbapenemase content, and β-lactamase content. Please refer to the corresponding spreadsheet file.**

**Table S3 legend:**

This table includes information on porin status, carbapenemase content, and β-lactamase content. For OmpC (OmpK36), certain annotations have been made based on previous studies. In 2019, Wong et al. demonstrated that a two-amino-acid insertion in OmpK36 resulted in increased carbapenem resistance (3). Additionally, Wong et al. (2022) identified a synonymous C-to-T mutation in OmpK36 associated with carbapenem resistance (4). Furthermore, Suelter and Hanson found that the OmpC of *E. coli* ST131 frequently exhibits down-regulation (11). Some of the *E. coli* strains included in this collection possess the same OmpC sequence as *E. coli* ST131 and were noted as they might also display down-regulation.
